# Classification of visual cortex plasticity phenotypes following treatment for amblyopia

**DOI:** 10.1101/554576

**Authors:** Justin L. Balsor, David G. Jones, Kathryn M. Murphy

## Abstract

Monocular deprivation (MD) during the critical period (CP) has enduring effects on visual acuity and the functioning of the visual cortex (V1). This experience-dependent plasticity has become a model for studying the mechanisms, especially glutamatergic and GABAergic receptors, that regulate amblyopia. Less is known, however, about treatment-induced changes to those receptors and if those changes differentiate treatments that support the recovery of acuity versus persistent acuity deficits. Here we use an animal model to explore the effects of 3 visual treatments started during the CP (n=24, 10 male and 14 female); binocular vision (BV) that promotes good acuity versus reverse occlusion (RO) and binocular deprivation (BD) that causes persistent acuity deficits. We measured recovery of a collection of glutamatergic and GABAergic receptor subunits in V1 and modeled recovery of kinetics for NMDAR and GABA_A_R. There was a complex pattern of protein changes that prompted us to develop an unbiased data-driven approach for these high-dimensional data analyses to identify plasticity features and construct plasticity phenotypes. Cluster analysis of the plasticity phenotypes suggests that BV supports adaptive plasticity while RO and BD promote a maladaptive pattern. The RO plasticity phenotype appeared more similar to adults with high expression of GluA2 and the BD phenotypes were dominated by GABA_A_α1, highlighting that multiple plasticity phenotypes can underlie persistent poor acuity. After 2-4 days of BV the plasticity phenotypes resembled normals, but only one feature, the GluN2A:GluA2 balance, returned to normal levels. Perhaps, balancing Hebbian (GluN2A) and homeostatic (GluA2) mechanisms is necessary for the recovery of vision.

## Introduction

Since the earliest demonstrations that monocular deprivation (MD) during a critical period (CP) causes ocular dominance plasticity and acuity loss[1]-[3] this model has been used to deepen our understanding of the neural changes associated with amblyopia. There have been fewer studies, however, about cortical changes associated with the acuity deficits that often persist after treatment for amblyopia[4]-[8]. Here we use an animal model to classify the expression patterns (phenotypes) of a collection of synaptic proteins that regulate experience-dependent plasticity and explored if treatments that promote good versus poor acuity reinstate CP-like plasticity phenotypes in visual cortex (V1).

Many animal studies have highlighted the roles of glutamatergic and GABAergic mechanisms for regulating plasticity during the CP[9]-[15]. For example, the subunit composition of AMPA, NMDA, and GABA_A_ receptors regulate the bidirectional nature of ocular dominance plasticity[16]-[21]. Some of the changes caused by MD include delaying the maturational shift to more GluN2A-containing NMDARs[22], [23], and accelerating the expression of GABA_A_α1-containing GABA_A_Rs[20], [23]. Together those changes likely decrease signal efficacy and dysregulate the spike-timing dependent plasticity that drives long-term depression (LTD) and weakens deprived eye response[24]. Furthermore, silencing activity engages homeostatic mechanisms that scale the responsiveness of V1 neurons by inserting GluA2-containing AMPAR into the synapse[25]. Importantly, many of the receptor changes have been linked with specific acuity deficits[26], [27] suggesting that visual outcomes may reflect changes to a collection of glutamatergic and GABAergic receptor subunits that together represent a plasticity phenotype for V1.

Animal studies of amblyopia have also identified treatments that promote good versus poor recovery of acuity after MD. For example, reverse occlusion (RO) gives a competitive advantage to the deprived eye that promotes an ocular dominance shift but the acuity recovered by the deprived-eye is transient, and can be lost within hours of introducing binocular vision[6]-[8]. Similarly, closing both eyes after MD to test a form of binocular deprivation therapy (BD) leads to poor acuity in both eyes that does not recover even after months of binocular vision[28]. In contrast, just opening the deprived eye to give binocular vision (BV) after MD appears to engage cooperative plasticity that promotes both physiological recovery[29] and long-lasting visual recovery in both eyes[27].

Here we quantified expression of glutamatergic and GABAergic receptor subunits in V1 of animals reared with MD and then treated to promote either good visual recovery (BV) or persistent bilateral amblyopia (RO, BD). Next, we developed an unbiased high-dimensional analysis approach to identify plasticity features in the pattern of subunit expression and to construct plasticity phenotypes. Finally, we used cluster analysis to classify plasticity phenotypes associated with good versus poor acuity and analyzed those to determine which features suggest the recovery of adaptive versus maladaptive plasticity mechanisms.

## Materials & Methods

### Animals & Rearing Conditions

All experimental procedures were approved by the McMaster University Animal Research Ethics Board. We quantified the expression of 7 glutamatergic and GABAergic synaptic proteins in V1 of cats reared with MD from eye opening until 5 weeks of age and then given one of 3 treatments: RO for 18d, BD for 4d, or BV for either short-term (ST-BV, 1hr, 6hrs) or long-term (LT-BV, 1d, 2d or 4d) (n=7, 4 male and 3 female) (Figure 1). The lengths of RO and BD were selected because they have well documented and consistent visual changes that result in poor acuity in both eyes[7], [8], Murphy:1991td [30]. The BV periods were selected to match the lengths used previously to study rapid and dynamic changes caused by MD in both cat and mouse V1 [27], [31], [32]. Also, the short- and long-term BV groups were based on the data-driven analysis of protein expression described in detail below and that analysis placed the samples from ST-BV (1hr or 6hrs) versus LT-BV (1d, 2d or 4d) rearing conditions into separate clusters. The raw data collected previously[23] from animals reared with normal binocular vision until 2, 3, 4, 5, 6, 8, 12, 16, or 32 wks of age (n=9 animals, 2 male and 7 female), or MD from eye opening (6-11d) to 4, 5, 6, 9, or 32 wks (n=8 animals, 4 male and 4 female) were used for comparison.

**Figure 1.**
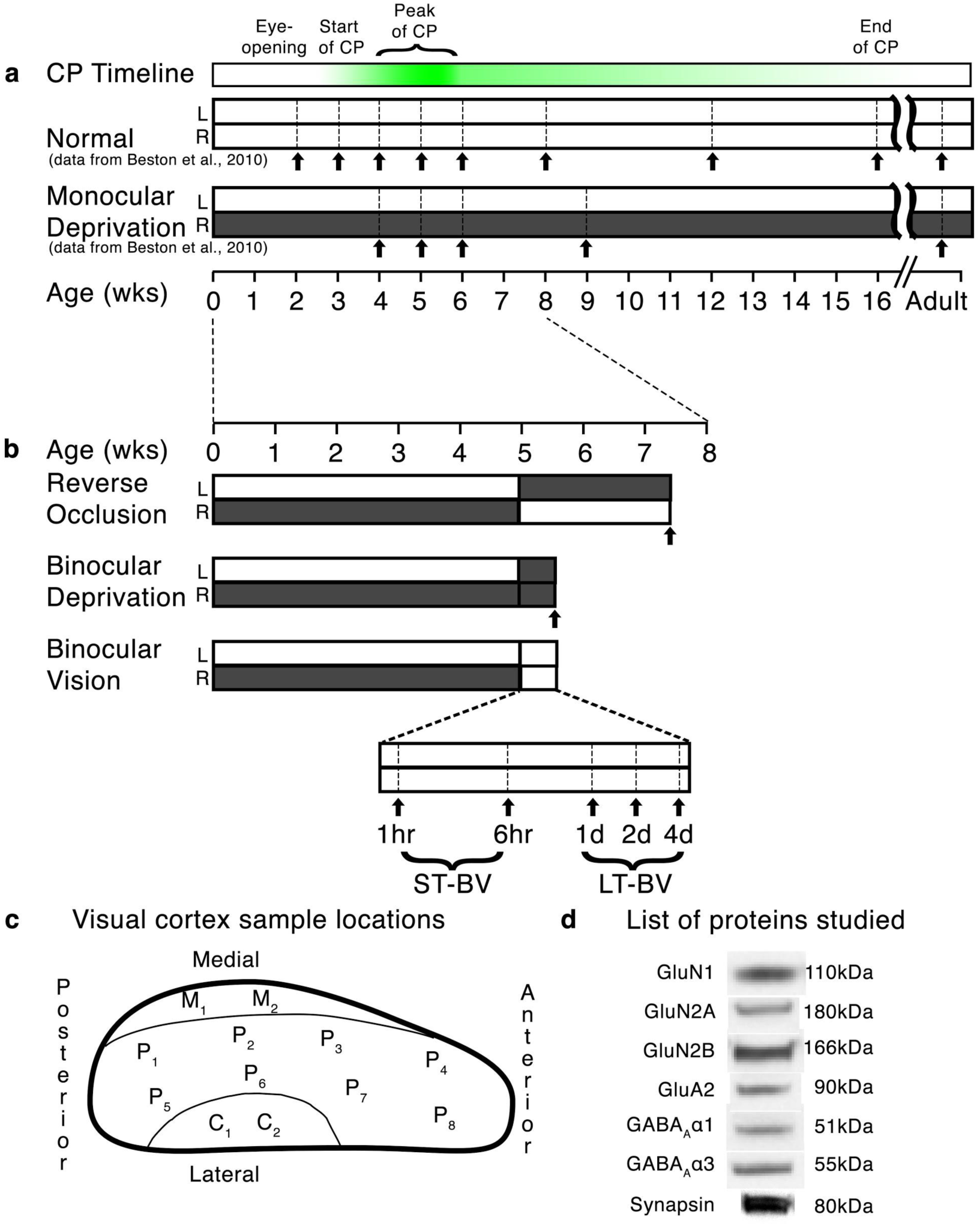
Study design diagram. Timelines for the rearing conditions used in this study. **a**. Normal visual experience and monocular deprivation (MD), **b**. treatment conditions (RO, BD, BV) after MD to 5wks. Filled bars indicate that an eye was closed. Black arrows indicate the age of animals used in the study. A timeline for the critical period (CP) in cat visual cortex[34] highlights the peak of the CP between 4-6 weeks of age. **c**. Map of the sampling regions in V1 representing central (C, n=2), peripheral (P, n=8), and monocular (M, n=2) visual fields. **d**. Representative bands from the Western blots for the 7 proteins quantified in the study and the molecular weights (kDa).

MD was started at the time of eye-opening by suturing together the eyelid margins of one eye (5-0 Coated VICRYL Ethicon P-3) using surgical procedures described previously[8]. Sutures were inspected daily to ensure the eyelids remained closed. At 5 weeks of age, the period of MD was stopped and either BV was started by carefully parting the fused eyelid margins, RO was started by opening the closed eye and closing the open eye or BD was started by closing the open eye. All of these surgical procedures were done using gaseous anesthesia (isoflurane, 1.5-5%, in oxygen) and aseptic surgical techniques.

At the end of the rearing condition animals were euthanized using sodium pentobarbital injection (165mg/kg, IV), and transcardially perfused with cold 0.1M phosphate buffered saline (PBS) (4°C; 80-100 ml/min) until the circulating fluid ran clear. The brain was removed from the skull and placed in cold PBS. A number of tissue samples (2 mm × 2 mm) were taken from the regions of V1 representing the central (C), peripheral (P) and monocular (M) visual fields (Figure 1c). Each tissue sample was placed in a cold microcentrifuge tube, flash frozen on dry ice, and stored in a −80°C freezer.

### Synaptoneurosome preparation

Synaptoneurosomes were prepared according to a subcellular fractionation protocol[16], [33]. The tissue samples were suspended in 1 ml of cold homogenization buffer (10 mM HEPES, 1 mM EDTA, 2 mM EGTA, 0.5 mM DTT, 10 mg/l leupeptin, 50 mg/l soybean trypsin inhibitor, 100 nM microcystin and 0.1mM PMSF), and homogenized in a glass-glass Dounce tissue homogenizer (Kontes, Vineland, NJ, USA). Homogenized tissue was passed through a 5µm-pore hydrophobic mesh filter (Millipore, Billerica, MA), centrifuged at low-speed (1,000xg) for 20 min, the supernatant was discarded, and the pellet was re-suspended in 1ml cold homogenization buffer. The sample was centrifuged for 10 min (1000x*g*), the supernatant was discarded, and the pellet was re-suspended in 100µl boiling 1% sodium-dodecyl-sulfate (SDS). Samples were heated for 10 min and then stored at −80°C.

Total protein concentrations were determined for each sample and a set of protein standards using the bicinchoninic acid (BCA) assay (Pierce, Rockford, IL, USA). A linear function was fit to the observed absorbance values of the protein standards relative to their expected protein concentrations. If the fit was less than R^2^=0.99, the assay was re-done. The slope and the offset of the linear function were used to determine the protein concentration of each sample and then the samples were diluted to 1 µg/µl with sample (M260 Next Gel Sample loading buffer 4x, Amresco) and Laemmli buffer (Cayman Chemical). A control sample was made by combining a small amount from each sample to create an average sample that was run on every gel. Each sample was run twice in the experiment.

### Immunoblotting

Synaptoneurosome samples and a protein ladder were separated on 4-20% SDS-PAGE gels (Pierce, Rockford, IL) and transferred to polyvinyldenine fluoride (PVDF) membranes (Millipore, Billerica, MA). The blots were blocked in PBS containing 0.05% Triton-x (Sigma, St. Louis, MO) (PBS-T) and 5% skim milk (wt/vol) for 1 hour. Blots were then incubated overnight at 4°C with constant agitation in one of the 7 primary antibodies (Table 1), and washed with PBS-T (Sigma, St. Louis, MO) (3 × 10 min).

**Table1:** List of primary antibody concentrations.

| **Antibody** | **Concentration** | **Company** | **Lot Number** | **Location** | **RRID** |
| --- | --- | --- | --- | --- | --- |
| anti-GluN1 | 1:2000 | BD Biosciences<br>Pharmingen | 556308 | San Diego, CA | RRID:AB_396353 |
| anti-GluN2A | 1:2000 | Millipore Sigma | 24826 | Burlington, MA | RRID: AB_95169 |
| anti-GluN2B | 1:2000 | Millipore Sigma | 28629 | Burlington, MA | RRID: AB_2112925 |
| anti-GluA2 | 1:1000 | Thermo Fisher |  | Waltham, MA | RRID: AB_2533058 |
| anti-GABAα1 | 1:500 | Santa Cruz<br>Biotechnology | L3102 | Santa Cruz, CA | |
| anti-GABAα3 | 1:2000 | Millipore Sigma | | Burlington, MA | |
| anti-Synapsin | 1:2000 | Thermo Fisher |  | Waltham, MA |  |

The appropriate secondary antibody conjugated to horseradish peroxidase (HRP) (1:2000; Cedarlane laboratories LTD, Hornby, ON) was applied to membranes for 1 hour at room temperature, then blots were washed in PBS (3 × 10 min). Bands were visualized using enhanced chemiluminescence (Amersham, Pharmacia Biotech, Piscataway, NJ) and exposed to autoradiographic film (X-Omat, Kodak, Rochester, NY). After each exposure blots were stripped (Blot Restore Membrane Rejuvenation kit (Chemicon International, Temecula, CA, USA)) and probed with the next antibody so each blot was probed for all 7 antibodies (Figure 1d).

### Analysis of Protein Expression

The autoradiographic film and an optical density wedge (Oriel Corporation, Baltimore, MD) were scanned (16 bit, AFGA Arcus II, Agfa, Germany), and the bands were identified based on molecular weight. The bands were quantified using densitometry and the integrated grey-level of the band was converted into optical density units (OD) using custom software (MATLAB, The Mathworks, Inc., Natick, Massachusetts). The background density between the lanes was subtracted from each band and the density of each sample was normalized relative to the control sample run on each gel (sample band density/control band density).

The data were normalized relative to the average expression of the 5wk normal cases. Table 2 summarizes the number of tissue samples and replication of runs for the 5wk Normal, 5wk MD and recovery conditions across the 3 regions of V1, and 7 proteins that were studied. Descriptions of the expression for the individual proteins in each of the conditions can be found in [35]. Those univariate comparisons confirmed the complex nature of these data and led us to develop and implement the data analysis workflow that is summarized in Figure 2.

**Figure 2.**
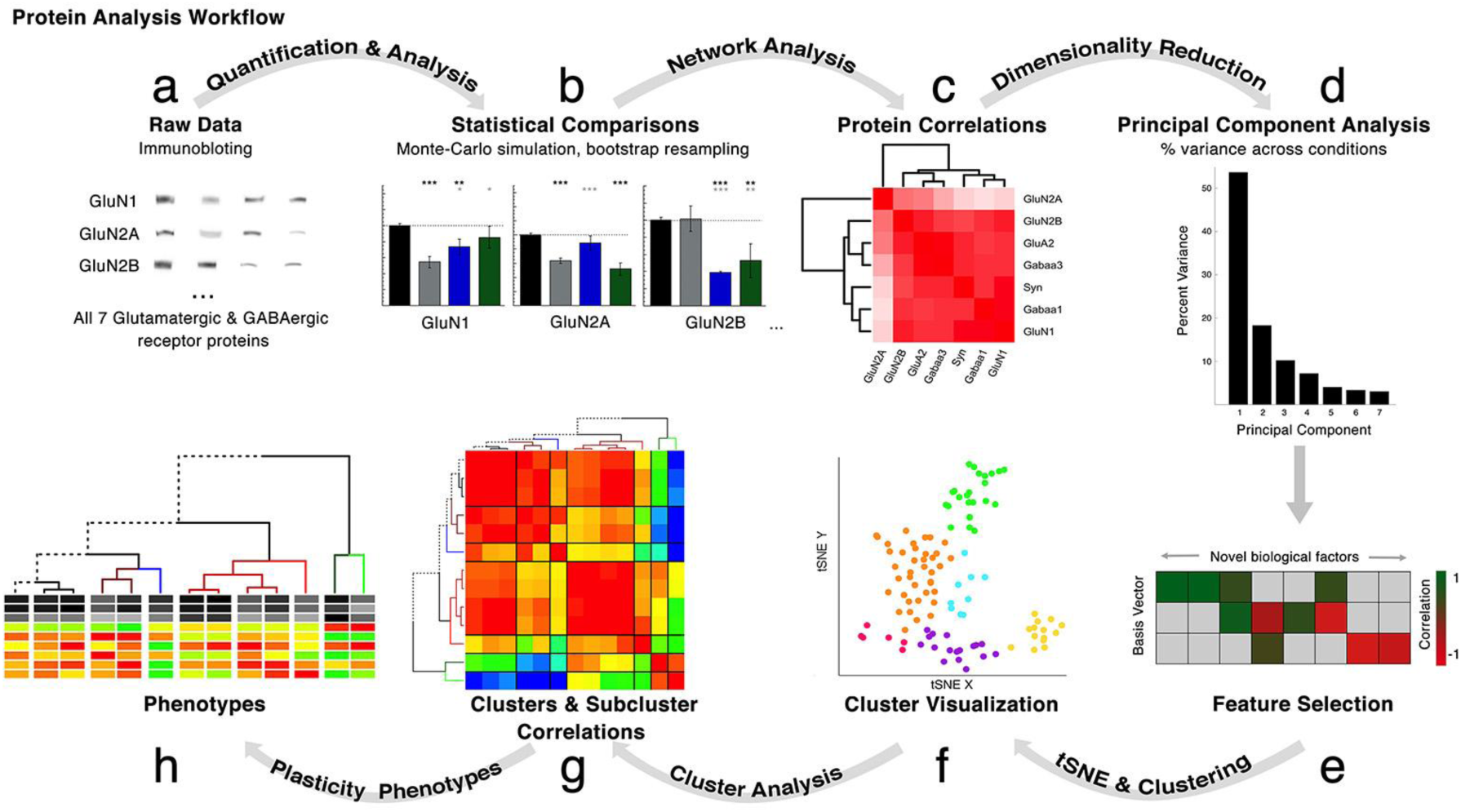
Analysis workflow. The analysis workflow for data in the study. **a**. Immunoblots were quantified using densitometry, **b**. Comparisons among rearing conditions were made [35]. **c**. Pairwise correlations were calculated for the 7 proteins for each rearing condition. **d**. Next, a series of steps were done beginning with dimension reduction (PCA), **e**. Feature selection, **f**. Cluster visualization based on the features (tSNE), **g**. Correlation between features or the clusters and subclusters, **h**. Construction and visualization of the plasticity phenotypes for each subcluster.

**Table 2:** The number of animals, cortical tissue pieces, and WB measurements for each condition and V1 region. Rows summarize the number of runs from the Central (C), Peripheral (P), and Monocular (M) regions of V1 within a rearing condition. The columns list each of the 7 proteins analyzed using Western blotting. Column sums detail the number of runs across rearing conditions and cortical areas. The information for Normal animals is in Table 2-1, and for MD animals is in Table 2-2.

| Number of Western Blot measurements after 2 replications |  |  |  |  |  |  |  |  |  |  |
| --- | --- | --- | --- | --- | --- | --- | --- | --- | --- | --- |
| Condition | Number of Animals | Region | Number of Cortical Pieces | GluN1 | GluN2A | GluN2B | GABAα1 | GABAα3 | GluA2 | Synapsin |
| Normal (5wks) | 1 | C | 2 | 4 | 4 | 4 | 4 | 4 | 4 | 4 |
|  |  | P | 8 | 16 | 16 | 16 | 15 | 16 | 16 | 16 |
|  |  | M | 2 | 4 | 4 | 4 | 4 | 4 | 4 | 4 |
| MD (5wks) | 2 | C | 3 | 6 | 6 | 6 | 6 | 6 | 6 | 4 |
|  |  | P | 9 | 18 | 18 | 18 | 18 | 18 | 18 | 12 |
|  |  | M | 3 | 5 | 5 | 5 | 5 | 5 | 5 | 4 |
| RO (18d) | 1 | C | 2 | 4 | 4 | 4 | 4 | 4 | 4 | 4 |
|  |  | P | 8 | 19 | 19 | 19 | 19 | 19 | 14 | 14 |
|  |  | M | 2 | 3 | 3 | 3 | 3 | 3 | 2 | 2 |
| BD (4d) | 1 | C | 3 | 6 | 6 | 5 | 6 | 5 | 6 | 5 |
|  |  | P | 9 | 18 | 18 | 17 | 16 | 18 | 18 | 17 |
|  |  | M | 2 | 4 | 4 | 3 | 4 | 4 | 4 | 3 |
| ST-BV (1hr, 6hr) | 2 | C | 4 | 8 | 8 | 8 | 8 | 8 | 8 | 8 |
|  |  | P | 16 | 32 | 32 | 32 | 32 | 32 | 32 | 32 |
|  |  | M | 4 | 8 | 8 | 8 | 8 | 8 | 8 | 7 |
| LT-BV (1d, 2d, 4d) | 3 | C | 6 | 12 | 10 | 12 | 11 | 12 | 12 | 10 |
|  |  | P | 24 | 43 | 40 | 43 | 43 | 42 | 43 | 40 |
|  |  | M | 6 | 12 | 12 | 12 | 12 | 12 | 12 | 12 |
| SUM |  |  |  | 222 | 217 | 219 | 218 | 220 | 216 | 198 |

### Protein Network Analysis

A network analysis of protein expression was done for each rearing condition by calculating the pairwise Pearson’s R correlations among the 7 proteins using the *rcorr* function in the Hmisc package in R[36]. The networks were visualized as correlation matrices (*heatmap2* function in *gplots*[37]) and the proteins were ordered using the dendextend[38] and seriation[39] packages to place proteins with similar patterns of correlations nearby in the dendrogram. Significant correlations were identified using Bonferroni corrected p-values and indicated by asterisks on the cell in the correlation matrix.

### Principal component analysis

We used principal component analysis (PCA) to reduce the dimensionality of the data, identify potential biological features, and create plasticity phenotypes. We applied PCA following procedures we used previously[23], [40], [41] and included data from all of the normal animals and MDs as well as the 3 recovery conditions. We assembled protein expression for GluA2, GluN1, GluN2A, GluN2B, GABA_A_α1, GABA_A_α3, and Synapsin into an *mxn* matrix. The *m* columns represented the 7 proteins and the *n* rows were the average protein expression for each of the 12-14 samples from an animal. For a few of the rows data was missing from a single cell and so those samples were omitted for a total of *n*=279 rows in the matrix and 1,953 observations.

The data were centered by subtracting the mean column vector and applying a singular value decomposition (SVD) to calculate the principal components (R Studio). SVD represents the expression of all 7 proteins within a single tissue sample as a vector in high dimensional space and the PCA identifies variance captured by each dimension in that “protein expression space”. The first 3 dimensions accounted for 82% of the total variance and were used for the next analyses.

We plotted the basis vectors for the first 3 dimensions (Dim) and used the weight, quality (cos^2^) and directionality of each protein, as well as known protein interactions, to help identify potential biological features accounting for the variance. We identified 9 potential features, calculated those features for each sample and correlated each feature with Dim1, Dim2 and Dim3 to create a correlation matrix (see results). The p-values for the correlations were Bonferroni corrected and significant correlations were used to identify features that would be part of the plasticity phenotype.

Eight of the features were significantly correlated with at least one of the first 3 dimensions. A measure associated with the E:I balance, was not significantly correlated with the dimensions and so it was not included in the tSNE or cluster analysis. The E:I measure, however, was used for analyzing the composition of the clusters and as a component of the plasticity phenotype because of the importance of the E:I balance for experience-dependent plasticity.

### tSNE dimension reduction and cluster analysis

The average expression for the 8 features (Table 3) was compiled into an *m*x*n* matrix, with *m* columns (*m*=8) representing the significant features and *n* rows representing each sample from the 3 V1 regions (central, peripheral, monocular) for 5wk Normal, 5wk MD, RO, BD and BV animals (n=109). t-distributed stochastic neighbor embedding (t-SNE) was used to reduce this matrix to 2-dimensions (2D). tSNE was implemented in R[42] and the tSNE output was sorted using k-means to assign each sample to a cluster. To determine the optimal number of clusters (k) we calculated the within-groups sum of squares for increasing values of k, fit a single-exponential tau decay function to those data, found the “elbow point” at 4τ which was 6, and used that as the optimal number of clusters. The clusters were visualized by color-coding the dots in the tSNE plot and the composition of the clusters was analyzed.

**Table 3:**
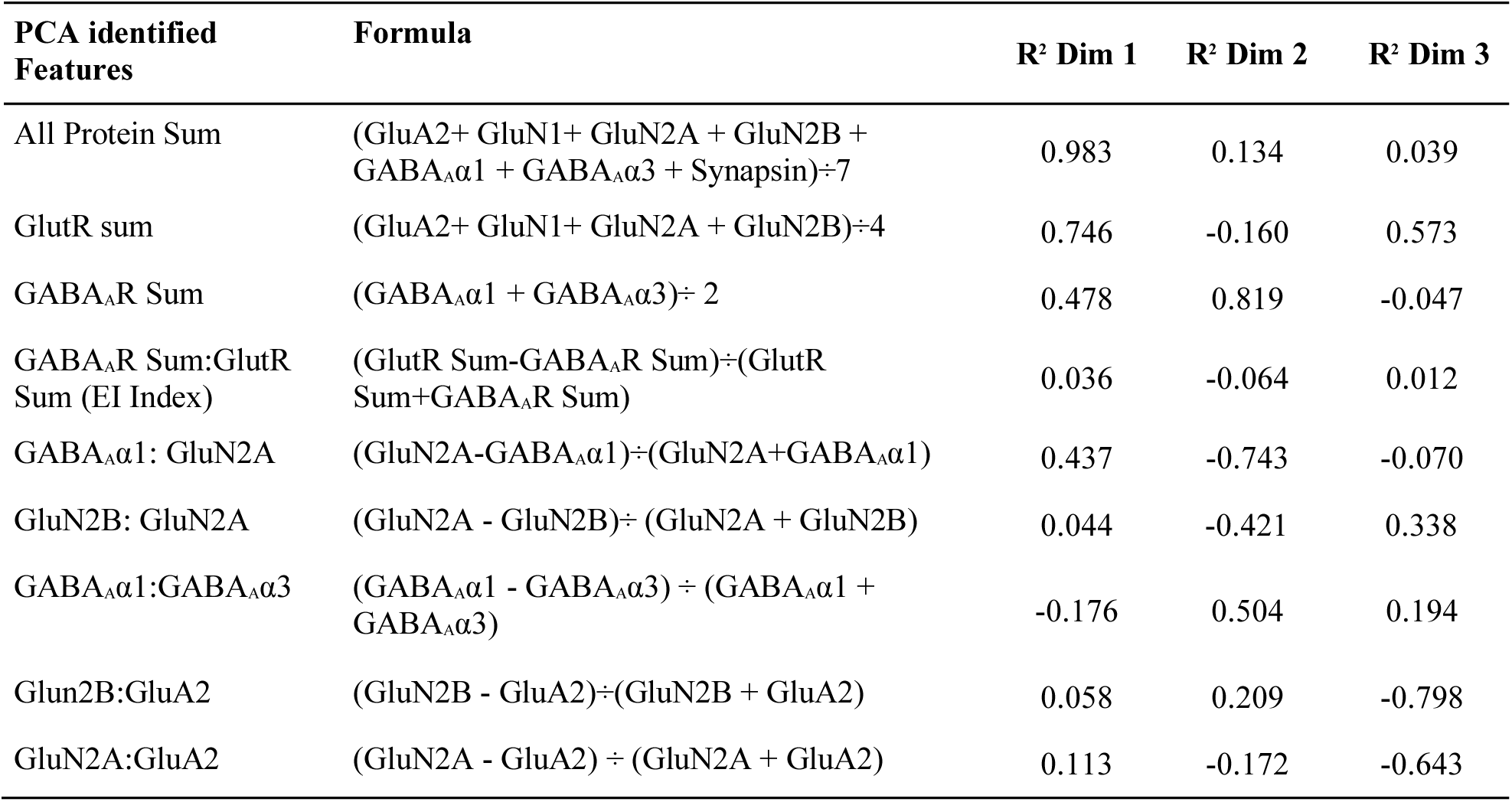
Formulas and Pearson’s R correlation between the features and principal components. The formulas for PCA identified features, including protein sums (Figure 5) and receptor indices (Figure 6), along with corresponding correlation (R^2^) values for each of the first 3 principal components. The GluN1:GluA2 and GABA_A_R Sum:GlutR Sum were not significantly correlated with any of these 3 components.

To facilitate analysis of the tSNE clusters we grouped the BV cases into short-term BV (1hr & 6hr) (ST-BV) or long-term BV (1d, 2d, and 4d) (LT-BV), color-coded the samples by rearing condition and used different symbols to indicate the V1 region. For each cluster we annotated the composition based on the rearing condition of the samples to create ‘subclusters’ (e.g. LT-BV 1) that were used for the next analyses.

We evaluated the similarity/dissimilarity among the subclusters by calculating the pairwise correlations (Pearson’s R) between subclusters using the features identified by the PCA as input to the R package *rcorr*. The correlations were visualized in a matrix with the cells color-coded to indicate the strength of the correlation[37]. The order of the subcluster in the matrix was optimized using hierarchical clustering and a dendrogram was created based on the pattern of correlations (using dendextend and seriation packages in R) so that subclusters with strong correlations were nearby in the dendrogram.

### Visualization and comparison of plasticity phenotype

The features identified in the PCA analysis were used to indicate the plasticity phenotype for each of the subclusters. In addition to the 8 significant features, the E:I measure was included in the visualization of the plasticity phenotype. The features were color-coded using grey scale for the 3 protein sum features and a color gradient (red = −1, yellow = 0, green = +1) for the 6 protein indices. The plasticity phenotypes were displayed as a stack of color-coded bars with one bar for each feature. For the subclusters, the plasticity phenotypes were ordered by the dendrogram to facilitate comparison among subclusters that were similar versus dissimilar. We also calculated the plasticity phenotypes for the full complement of normally reared and MD animals and displayed those in a developmental sequence to facilitate age-related comparisons with the recovery subclusters. Finally, we did a bootstrap analysis to determine which features of the plasticity phenotypes were different from 5wk normals and used Bonferroni correction to adjust the significance for the multiple comparisons. This analysis was displayed 2 ways: first, each of the 9 feature bands for the dendrogram ordered subclusters was color-coded white if it was not different, red if it was greater, and blue if it was less than 5wk normals; second, boxplots were made to show the value for each of the 9 features and to identify the subclusters that were different from 5wk normals.

A detailed description of the network analysis, PCA, tSNE, clustering and phenotype construction, along with example code for each of these steps can be found in [43].

### Modeling population receptor decay kinetics for NMDARs and GABA_A_Rs

The subunit composition of NMDARs and GABA_A_Rs determines the decay kinetics of the receptor[44], [45] and so we used that information to build a model for the decay kinetics of a population of receptors for each of the rearing conditions. The decay kinetics of the most common NMDAR composition, triheteromeric receptors containing GluN2A and 2B is 50ms±3ms, while diheteromers NMDARs containing only GluN2B are slower (2B=333ms±17ms) and those containing only GluN2A are faster (2A=36ms±1ms) [44]. The decay kinetics of GABA_A_Rs with both α1 and α3 subunits is 49ms±23ms while receptors with only the α3 subunit are slower (129.0ms±54.0ms) and only α1 are faster (42.2ms±20.5ms) [45].

We used the relative amounts of GluN2A and 2B, or GABA_A_α1 and α3, as inputs to the model. Receptors containing GluN2A and 2B or GABA_A_α1 and α3 are the most common in the cortex, so the model maximized the number of these pairs which was limited by the subunit with less expression. The remaining proportion of the highly expressed subunit was divided by 2 and used to model the number of pairs for those receptors (2A:2A or 2B:2B; α1:α1 or α3:α3) in the population. The population decay kinetics were then modeled by inserting the relative amounts of the subunits into these formulas:

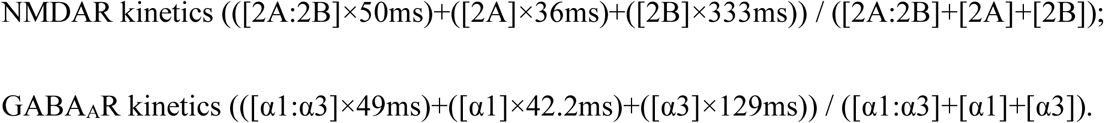

For example, a sample where GluN2A was 35% and 2B was 65% of the total NMDAR subunit population and would have population kinetics of 135ms.

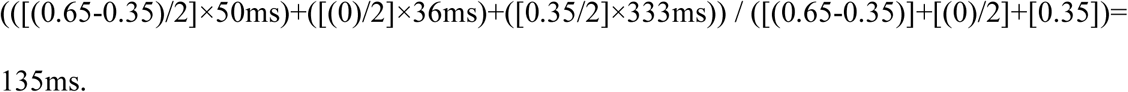

First, we plotted scattergrams of the average NMDAR and GABA_A_R decay kinetics for normal animals and each treatment condition. The development of decay kinetics for normal animals was described using an exponential decay function, while changes in kinetics with increasing lengths of BV were fit by either exponential decay or sigmoidal curves. Then we compared the relationship between NMDAR and GABA_A_R kinetics by plotting both on one graph.

### Statistical analyses

We used the bootstrap resampling method to compare the features because it is a conservative approach to analyzing small sample sizes when standard parametric or non-parametric statistical tests are not appropriate. Here bootstrapping was used to estimate the confidence intervals (CI) for each feature in the subcluster and a Monte Carlo simulation was run to determine if the 5wk Normal subcluster fell outside those CIs. The statistical software package R was used to simulate normal distributions with 1,000,000 points using the mean and standard deviation from the subcluster. Next, a Monte Carlo simulation randomly sampled with replacement from the simulated distribution *n* times, where *n* was the number of observations made from the normal subcluster. The resampling procedure was repeated 100,000 times to determine the 95%, 99% and 99.9% CIs. The subcluster feature was considered significantly different from normal (e.g., p<0.05, p<0.01 or p<0.001) if the feature mean fell outside these CIs. When a subcluster was significantly greater than normal (p<0.05) the boxplot was colored red, when it was less than normal (p<0.05) the boxplot was colored blue, and if not it was not different from normal (p>0.05), the boxplot was colored grey.

All of the bootstrap statistical comparisons for the plasticity features (Table 5-1 and 6-1) are presented in the Supplemental material.

The p-values for the Pearson’s correlations were calculated using the rcorr package[36], and the significance levels were adjusted using the Bonferroni correction for multiple comparisons. The Pearson’s Rs and p-values for the protein networks (Table 3-1), plasticity features with PCA dimensions (Table 4-1), and association between clusters (Tables 8-1, 8-2) are included in the Supplemental material.

We tested if recovery during BV followed either an exponential decay or sigmoidal pattern by fitting curves to the data using Kaleidagraph (Synergy Software, Reading PA). Significant curve fits were plotted on the graphs to describe the trajectory of recovery.

## Results

### Analyzing the pairwise similarity between protein expression profiles

First, we wanted to identify pairs of proteins with similar or opposing expression profiles and compare them among the rearing conditions. For each condition, we collapsed the data from the 3 regions of V1, calculated the matrix of pairwise correlations between the 7 proteins, ordered the protein correlations using a hierarchical dendrogram, and used 2D heatmaps to visualize the correlations (Figure 3). The order of proteins in the dendrogram indicated which ones had similar (e.g. on the same branch of the dendrogram) or different patterns of expression and the color of the cell illustrated the strength of the correlation. For 5wk normal animals (Figure 3a), there were strong positive correlations (red cells) among all proteins except GluN2A, which was weakly correlated and not clustered with the other proteins. A different pattern of correlations was found after MD (Figure 3b); here glutamatergic proteins were weakly, or even negatively correlated (blue cells) with GABA_A_α1, GABA_A_α3, and synapsin. These results suggest that MD drives a decoupling of these excitatory and inhibitory mechanisms. RO also separated glutamatergic and GABAergic proteins into different clusters at the first branch (Figure 3c); however, the correlations were weaker, suggesting that RO reduced the MD-driven decoupling of these mechanisms. After BD the correlation matrix had mostly positive correlations (Figure 3d) except for synapsin which was negatively correlated and not clustered with the other proteins. BV treatment highlighted the dynamic nature of this recovery (Figure 3e-i). Just 1hr of BV was enough to change the correlation matrix from the MD pattern, but even after 4d of BV the correlation matrix still appeared different from the normal 5wk pattern of correlations.

**Figure 3.**
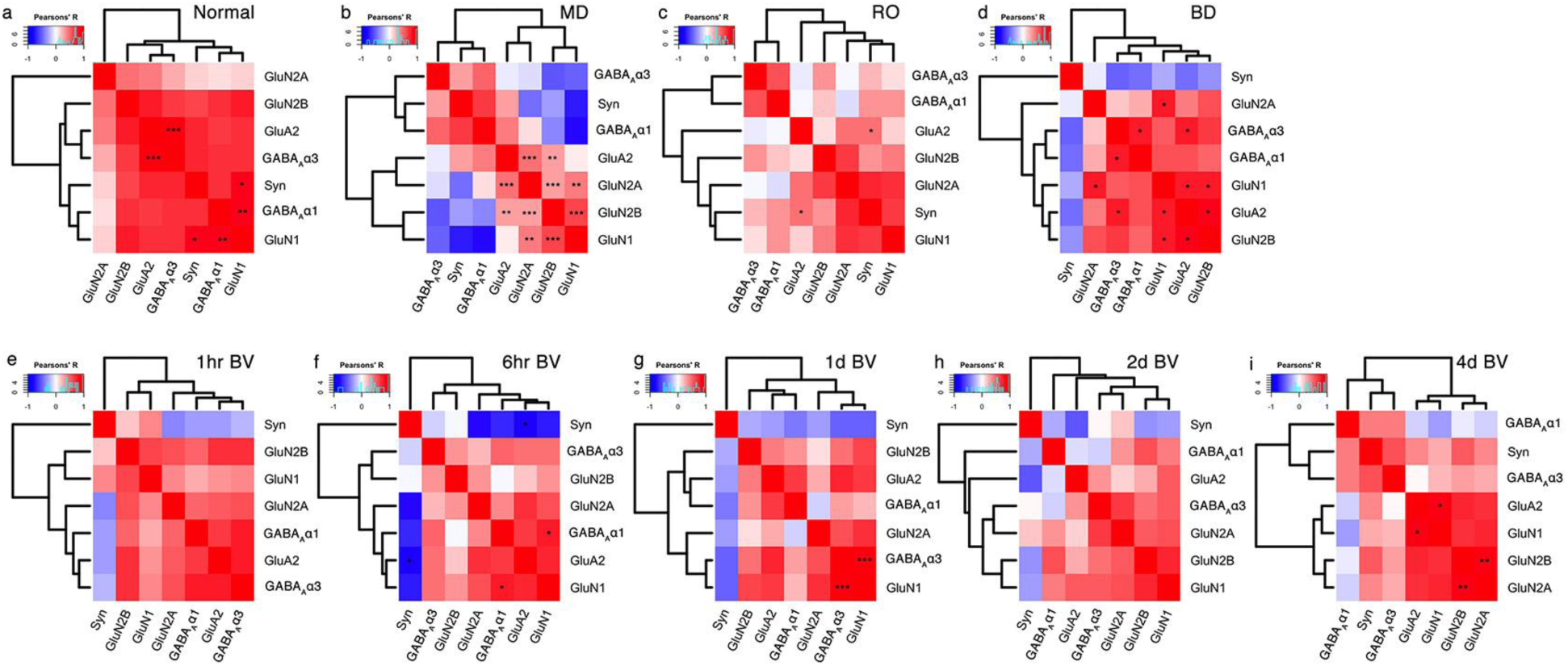
Visualizing pairwise correlations between proteins. Correlation matrices are plotted showing the strength (saturation) and direction (blue:negative; red:positive) of the pairwise Pearson’s R correlations between proteins for each condition **a**. 5wk Normal, **b**. 5wk MD, **c**. RO, **d**. BD, and **e-i**. BV. The order of proteins was determined using hierarchical clustering so proteins with stronger correlations were nearby in the matrix. Significant correlations are denoted by an asterisk (*p<0.05, **p<0.01, ***p<0.001). For table of Pearson’s R values and Bonferroni corrected p-values see Supplemental Table 3-1.

These matrices suggest different patterns of correlations depending on the condition, but this analysis treats each comparison with the same weighting and it is likely that some proteins contribute more than others to the variance in the data. To assess this, we used PCA to identify individual proteins and combinations of proteins that capture the variance in the data and may represent plasticity features reflecting differences among the treatment conditions.

### Using principal component analysis to reduce dimensionality and identify plasticity features

We used PCA to reduce the dimensionality, transform the data and find features that define the covariance among the proteins. An *m*x*n* matrix was made using protein expression, where the *m* columns were the 7 proteins and the *n* rows (109) were the tissue samples from all the animals and regions of V1 used in this study. This matrix was analyzed using singular value decomposition (SVD), and the first 3 dimensions explained most of the variance (82%) in the data (Dim1: 54%, Dim2: 18%, Dim3: 10%) (Figure 4a).

**Figure 4.**
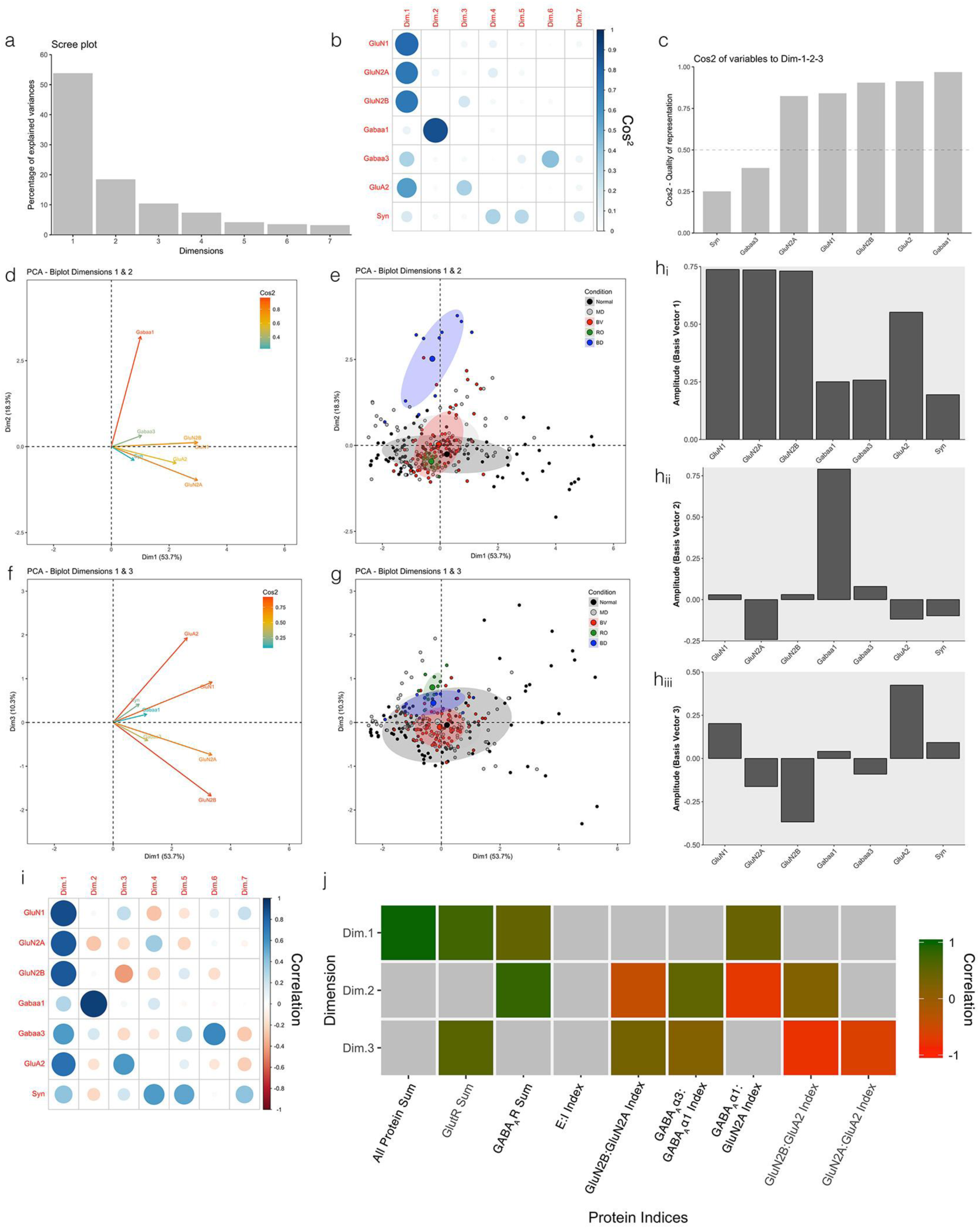
Identifying plasticity features using principal component analysis. **a**.The percentage of variance captured by each principal component by singular value decomposition (SVD) applied using all of the protein expression data. The first 3 principal components capture 54%, 18% and 10% of the variance, respectively, totalling >80% and thus representing the significant dimensions. **b.** The quality of the representation, cos^2^, for the proteins is plotted for each dimension (small/white: low cos^2^; large/blue: high cos^2^) **c**. The sum of cos^2^ values for the first 3 dimensions for each protein. **d,e.** Biplots of PCA dimensions 1&2 and **f,g.** 1&3. These plots show the vector for each protein (d,f) and the data (small dots) plus the average (large dots) for each condition with the best-fitting ellipse (e,g). **h.** The basis vectors for dimensions 1-3 showing the amplitude of each protein in the vector. **i.** The strength (circle size) and direction (blue-positive, red-negative) of the correlation (R^2^) between each protein and the PCA dimensions. **j.** Correlation between the plasticity features (columns) identified using the basis vectors (see Results) and then PCA dimensions 1-3. Filled cells are significant, Bonferroni corrected correlations (green = positive, red = negative). For table of Pearson’s R values and significant p-values for these associations see Supplemental Table 4-1.

To understand which proteins contributed to each dimension we addressed the quality of the representation for each protein using the cos^2^ metric and found that the glutamatergic proteins were well represented by Dim1, GABA_A_α1 by Dim2, and GluA2 and GluN2B by Dim3 but synapsin and GABA_A_α3 were weakly represented in the first 3 dimensions (Figure 4 b,c). Next, we compared the vectors for each protein (Figure 4 d,f) and the PCA space occupied by the rearing conditions (Figure 4 e,g). The protein vectors show that GluN1, GluN2A, GluN2B, and GluA2 extended along Dim1, GABA_A_α1 along Dim2, and GluA2 and GluN2B were in different directions along Dim3. The PCA space occupied by the conditions suggest some differences: BD was separated on Dim 2 in the same direction as GABA_A_α1, but the center of gravity for the other conditions overlapped the space occupied by normal samples.

The overlap among conditions raised the possibility that higher dimensions may separate the conditions. To begin to assess higher dimensional contributions we examined the basis vectors (Figure 4h) and the correlations between individual proteins and PCA dimensions (Figure 4i) to identify combinations of proteins that might reflect higher dimension features. For example, all proteins had positive amplitudes for the Dim1 basis vector (Figure 4h) and positive correlations with Dim1 (Figure 4i) suggested that protein sums may be higher dimensional features. In addition, weights for GluN2A and GABA_A_α1 on Dim2 were opposite, suggesting that when one protein increased the other decreased and this could be a novel feature of these data. Continuing with this approach we identified 9 putative plasticity features; protein sums (all protein sum, GlutR sum, GABA_A_R sum) or indices (GlutR:GABA_A_R, GluN2A:GluN2B, GABA_A_α1:GABA_A_α3, GluN2A:GABA_A_α1, GluA2:GluN2B, GluN2A:GluA2). All of the protein sums and 4 of the indices were features not analyzed with the univariate statistics; however, each had a strong biological basis in previous research. For example, the new indices paired the mature GluN2A with the mature GABA_A_α1 or GluA2 subunit, and GluN2B with GluA2 which is known to regulate the development of AMPARs[46]. Finally, we calculated the 9 features and determined if at least one of the first 3 dimensions was correlated with the features (Figure 4j). Only the GlutR:GABA_A_R balance was not correlated with any of the first 3 dimensions, but because those mechanisms are related to the E:I balance[47] we included that measure in the next analysis.

### Comparing plasticity features

We plotted the plasticity features and saw that the GlutR and GABA_A_R sums and indices identified various differences among the treatment conditions (Figures 5 & 6). There were, however, consistent changes after BV in the binocular regions with a loss of the total amount of GABA_A_R expression (44%±12) and a shift of the GlutR:GABA_A_R balance to favor GlutR (Figure 5d). The remaining indices in the feature list also identified differences (Figure 6) including the GABA_A_α1:GluN2A balance shifting to more GluN2A after BV (in binocular regions) but more GABA_A_α1 after BD. RO flipped the 2A:2B balance to favor more GluN2A as did BD in the central region. In contrast, BV shifted the 2A:2B balance towards normal CP levels in all of V1. The GABA_A_α1:GABA_A_α3 balance shifted towards the normal level after BV but strongly in favor of GABA_A_α3 after BD. The GluN2B:GluA2 balance shifted to substantially more GluA2 after RO while the GluN2A:GluA2 index shifted to more GluA2 outside the central region after RO and BD. Together, these features provide evidence of glutamatergic versus GABAergic differences among the treatment conditions.

**Figure 5.**
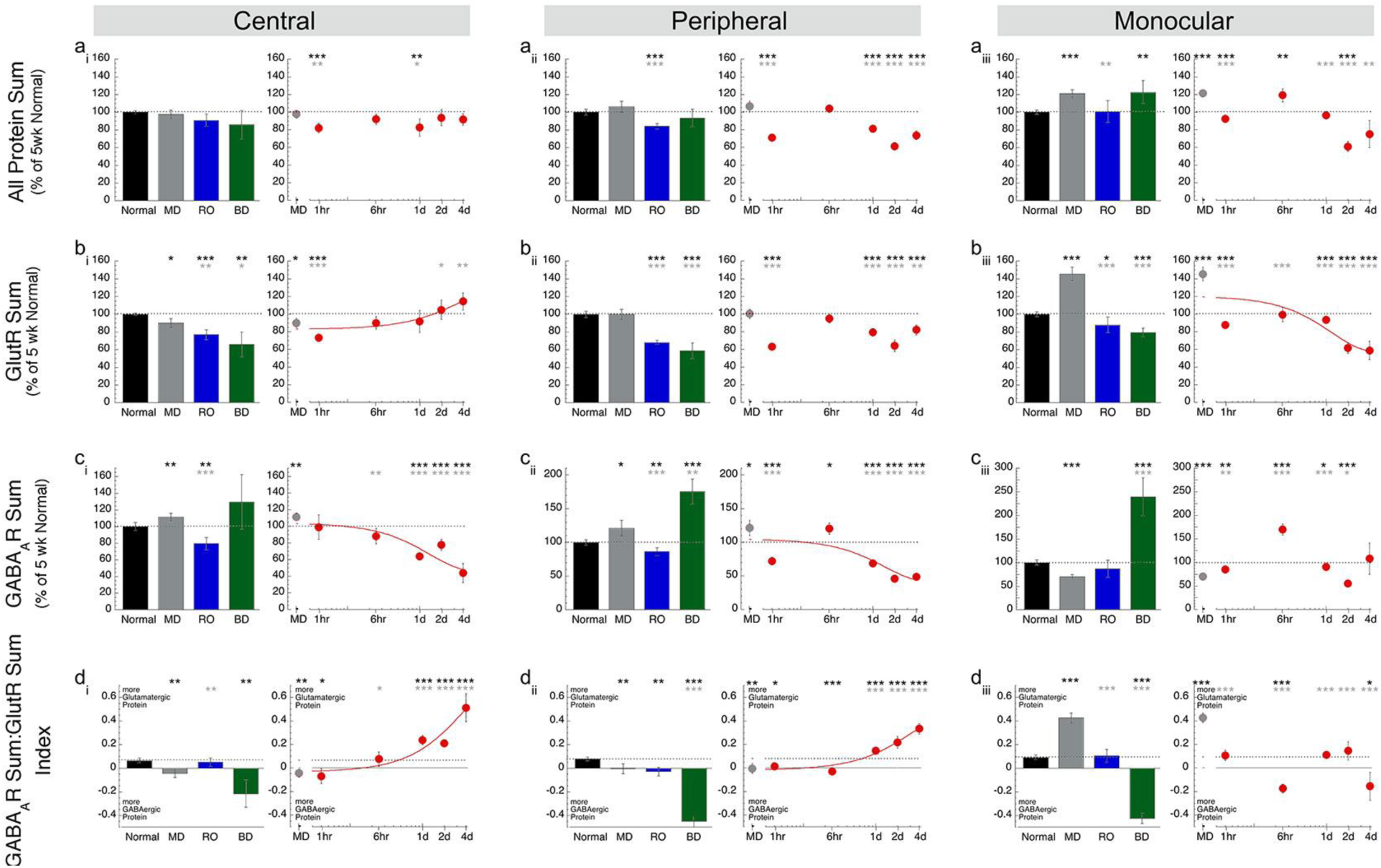
Expression of plasticity features for protein sums identified using principal component analysis. Histograms and scatterplots showing the protein sums and a new protein sum index (GABAR Sum:GlutR Sum-rows) that were identified using the PCA basis vectors (Fig. 4j) and plotted for each region of V1 (columns). For exact p-values, Pearson’s R, and equations for the curve-fits see Supplemental Table 5-1.

**Figure 6.**
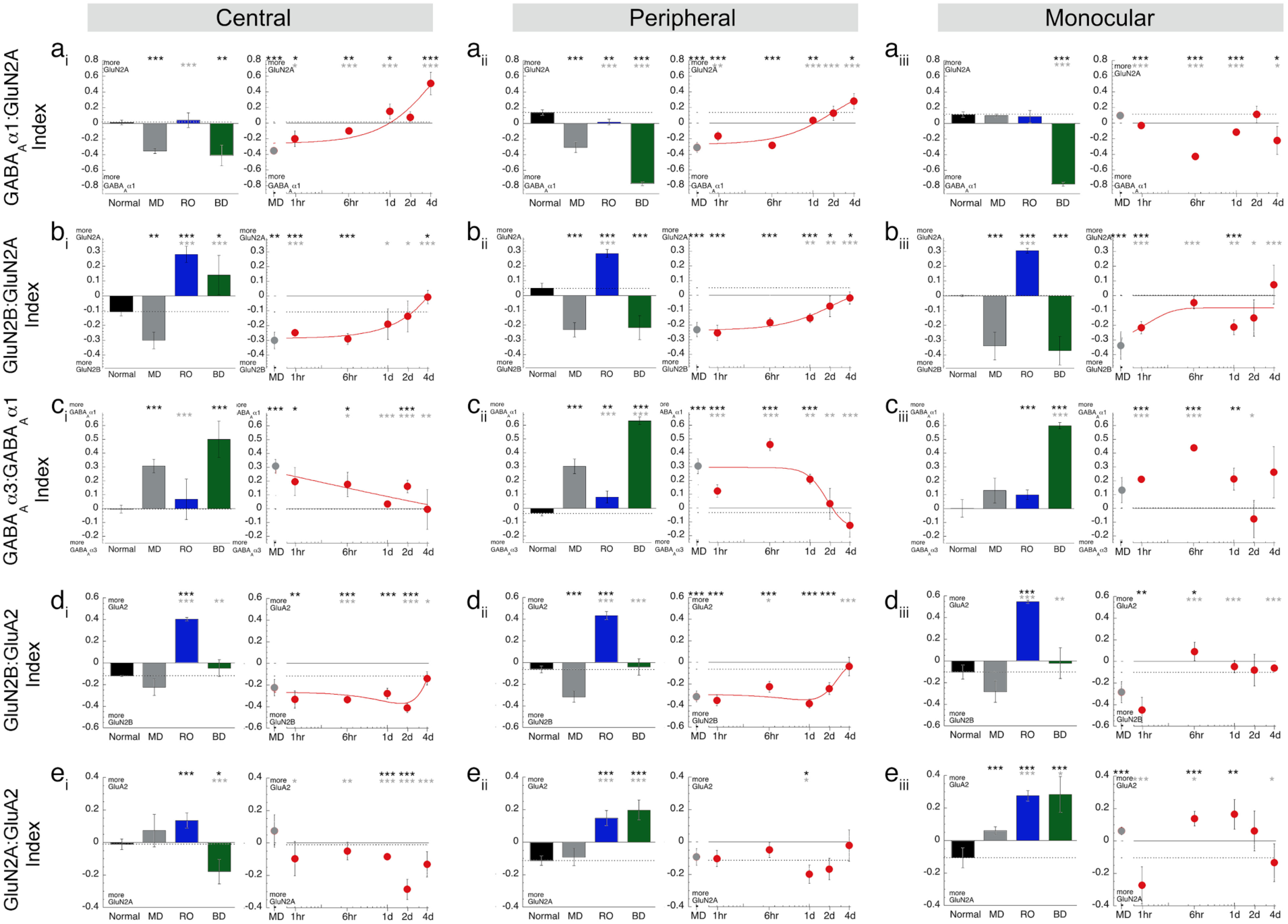
Expression of plasticity feature for indices identified using principal component analysis. Histograms and scatterplots showing the plasticity features (rows) that were identified using the PCA basis vectors (Fig. 4j) and plotted for each region of V1 (columns). The conventions are the same as in Figure 5. For exact p-values, Pearson’s R, and equations for the curve-fits see Supplemental Table 6-1.

### Using t-SNE to transform and visualize clustering in the pattern of plasticity features

We used t-SNE to transform the plasticity features and visualize them in 2D (Figure 7a), then k-means and the “elbow method” (Supplemental Figure 7-1) to identify the number of clusters. For these analyses, the BV samples were grouped into ST-BV (1-6hrs) and LT-BV (1-4d) groups, and the plasticity features were calculated for all samples from the 3 V1 regions.

**Figure 7.**
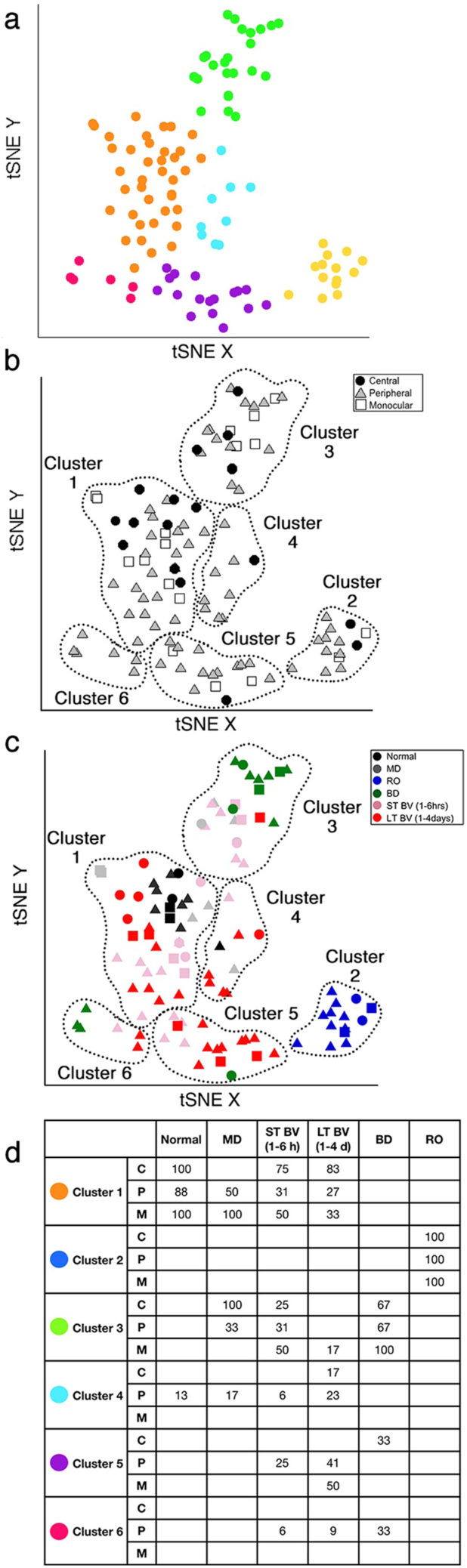
Clustering of samples with similar plasticity features identified using t-Distributed Stochastic Neighbor Embedding (t-SNE) and k-means clustering. **a.** Using tSNE to visualize clustering of samples (109 tissue samples from animals reared to 5wk normal, 5wk MD, RO, BD and BV) calculated from k-means analysis of the 8 plasticity features identified by PCA. The optimal number of clusters (k=6) was identified by measuring the within groups sum of squares at intervals between 2 and 9 clusters (Figure 7-1) **b.** The content of each cluster was visualized for the region (central, peripheral, monocular) **c**. or treatment condition. **d.** The table summarizes the percentage of samples for each region and condition in Cluster 1-6. For example, 100% of the samples from the central region of V1 in Normal animals were in Cluster 1 while 100% of the samples from all regions of RO were in Cluster 2. This information was used to annotate subclusters based on the cluster membership (1-6), rearing condition, and region of V1.

Six clusters were visualized with t-SNE (Figure 7) and the composition of the clusters was analyzed to determine the V1 regions and rearing conditions in each cluster. Cluster 1 was the largest with 39 samples (C=26%; P=54%; M=21%) and had the greatest number of samples from the central region (Figure 7 b,d). Cluster 3 also had samples from central, peripheral and monocular regions while clusters 4, 5, and 6 were dominated by peripheral samples with few or no central region samples. Thus, there was some clustering by V1 region, but more apparent clustering emerged when the samples were color-coded by rearing condition (Figure 7c, d). All but one of the normal samples were in cluster 1, all of the RO samples were in cluster 2, most of the BD samples were in cluster 3 with a few in cluster 6, and most of the MD samples were in clusters 1 or 3. The BV samples, however, were found in 5 of the clusters with the greatest number of BV central samples (83%) grouped with normals in cluster 1.

Further analysis of cluster 1 showed that the majority of LT-BV and ST-BV samples from the central region clustered with the normals (Figure 7d). Interestingly, some of the MD samples were also in cluster 1; however, those samples were from the peripheral and monocular regions which are known to be less affected by MD than the central region[48]. Together, these results show that the data are clustered and that the clustering was driven by both rearing condition and region of V1.

### Correlating plasticity features among subclusters

We annotated the samples in each cluster using the rearing condition and V1 region and used that information to identify 13 subclusters where at least one region per condition had n ≥ 2 and >20% of the samples in that cluster (Figure 7d, black font). A correlation matrix was calculated (Figure 8) to assess the similarity between subclusters (see Supplemental table 8-1 for R values and 8-2 for Bonferroni adjusted p values) and the order of the subclusters in the correlation matrix was optimized by hierarchical clustering so subclusters with similar patterns of features were nearby in the dendrogram. Bonferroni adjusted p value was used to determine the significant correlations (0.05/78=0.0006) (Figure 8). This analysis showed that 3 of the 4 LT-BV subclusters (LT-BV 1: R=0.98; LT-BV 5: R=0.98; LT-BV 4: R=0.96) and the MD 1_PM_ subcluster (R=0.98) were strongly correlated with normals. The other MD subcluster with central samples (MD 3_CP_) was on a separate branch of the dendrogram and was strongly correlated with the 3 ST-BV subclusters (ST-BV 1: R=0.98; ST-BV 3: R=0.99; ST-BV 5: R=0.98). The ST-BV subclusters were also correlated with normals (ST-BV 1: R=0.96; ST-BV 3: R=0.94; ST-BV 5: R=0.97), LT-BV 1 (ST-BV 1: R=0.98; ST-BV 3: R=0.94; ST-BV 5: R=0.98), and MD1 (ST-BV 1: R=0.98; ST-BV 3: R=0.94; ST-BV 5: R=0.99) but weaker correlations with LT-BV 4 (ST-BV 1: R=0.94; ST-BV 5: R=0.95) and no significant correlations with LT-BV 5. RO was correlated with normal (R=0.96) but only one of the LT-BV subclusters (LT-BV 5: R=0.96) and none of the ST-BV subclusters. The two BD subclusters were correlated (R=0.94) but none of the other correlations were significant. The pattern of strong correlations in this matrix and the resulting dendrogram suggested that the subclusters might form 4 groups that have similar plasticity features (1: normal, LT-BV, MD_P or M_; 2: RO; 3: ST-BV, MD_C_; 4: BD).

**Figure 8.**
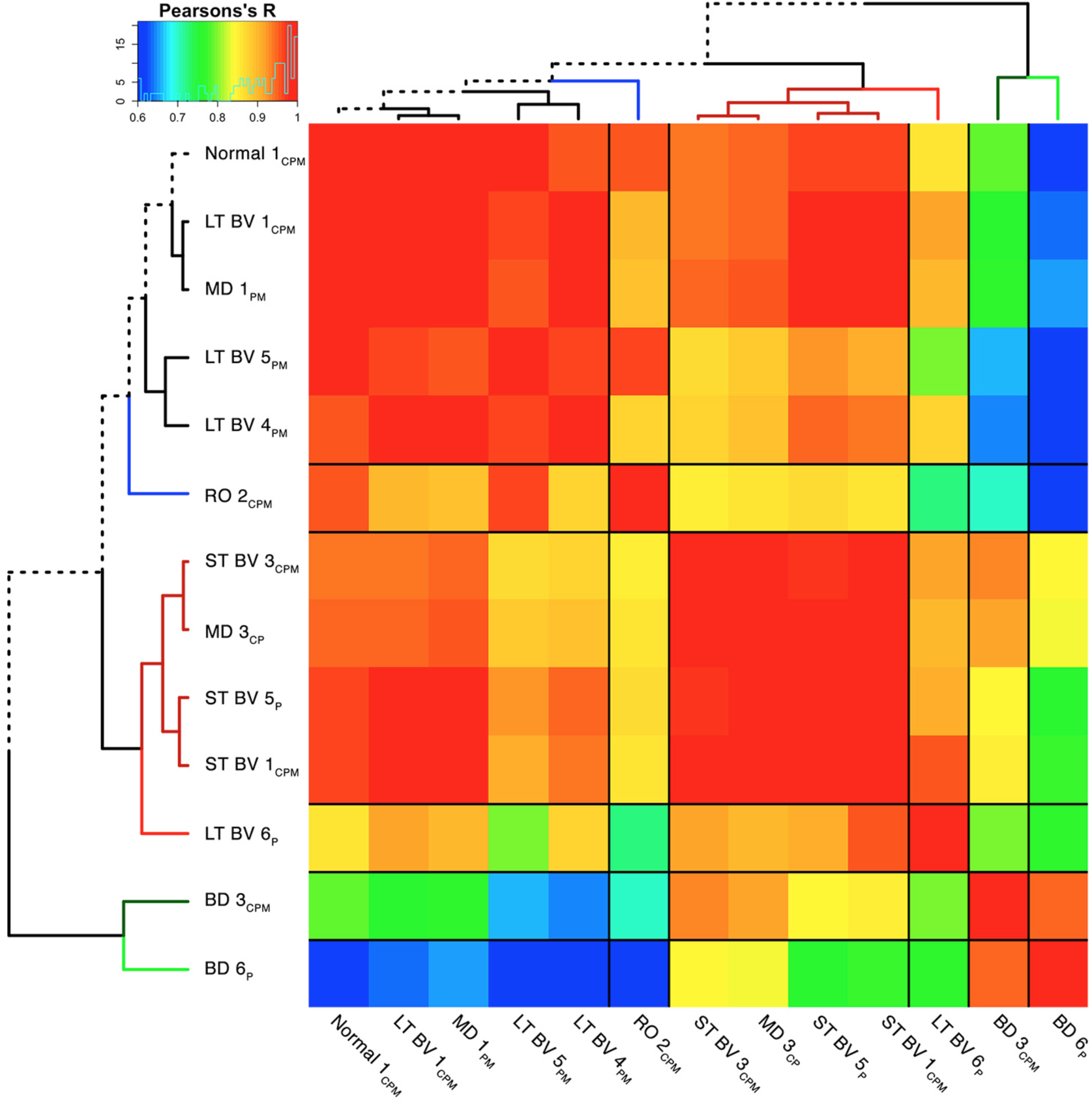
Visualizing pairwise correlations between treatment subclusters. The matrix is showing the strength (0.6=blue; 1=red) of correlation between the subclusters identified in Fig. 7d and annotated here using the rearing condition, cluster (1-6), and region of V1. Hierarchical clustering was used to order the data so that subclusters with strong correlations were nearby in the matrix. The subclusters formed 5 groups using the height of the dendrogram that is denoted by a change in the color of the dendrogram. The dotted black line in the dendrogram highlights the path to the normal subcluster. The black lines in the matrix identify the 5 groupings of the subclusters. For exact Bonferroni corrected p-values see Supplemental Table 8-1 and for Pearson’s R values see Supplemental Table 8-2.

### Constructing plasticity phenotypes and comparing among the subclusters

To compare the pattern of plasticity features among the subclusters we visualized the average for each feature as a color-coded horizontal band, stacked the bands to illustrate the pattern that we called the plasticity phenotype (Figure 9a) and ordered the phenotypes using the same dendrogram as the correlation matrix (Figure 9b). In addition, we visualized the plasticity phenotypes for normal development and MD (using the data from [23]) to compare the treatment subclusters with a broad range of ages that had developed with either normal or abnormal visual experience (Figure 9 c,d).

**Figure 9.**
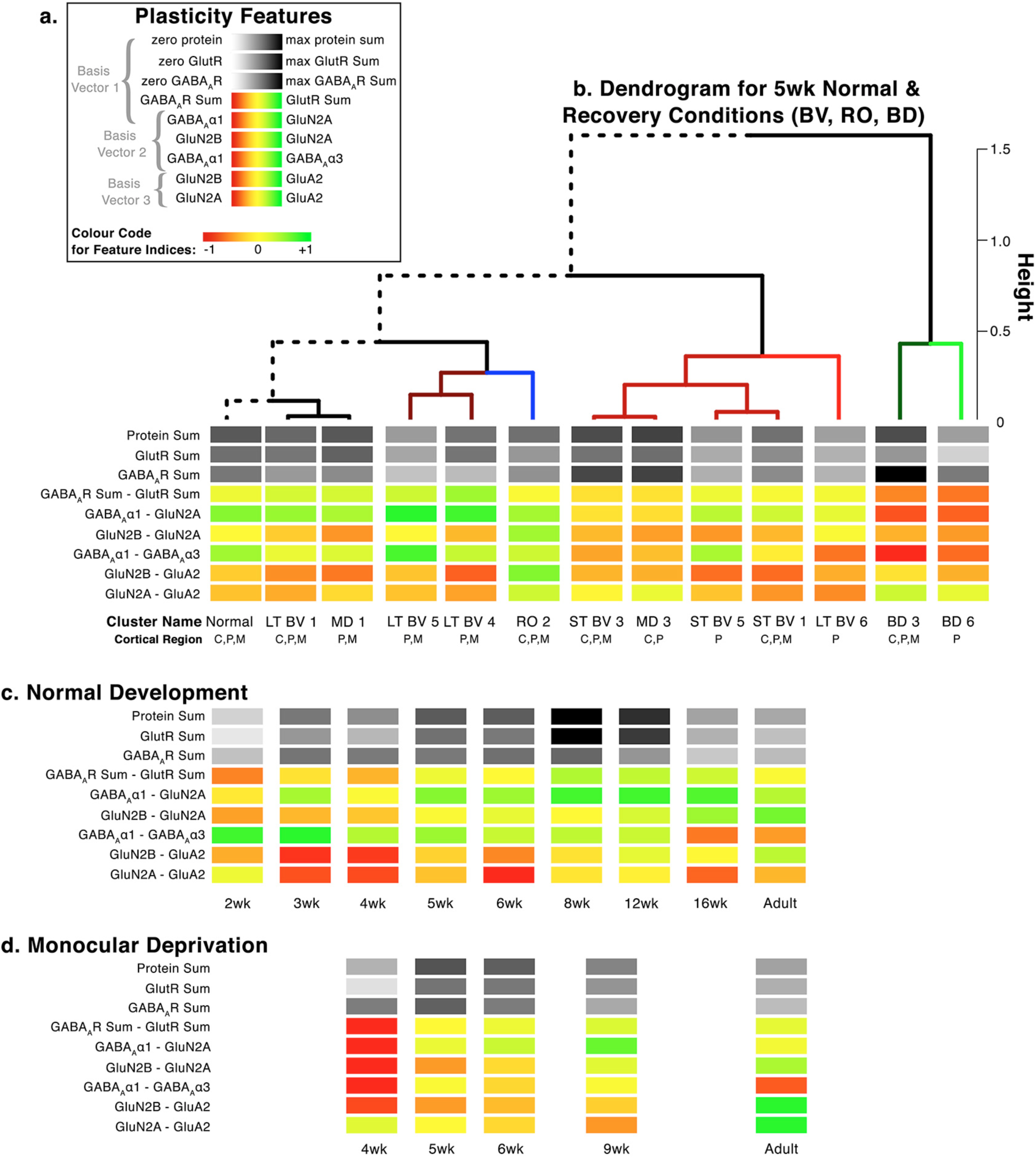
Visualizing the plasticity features and phenotypes for each subclusters. **a.** We visualized of the plasticity features as a stack of color-coded horizontal bars that together comprise the plasticity phenotype. The 3 grey scale bars represent the protein sums and the 6 red-green color-coded bars represent the protein indices identified by the PCA. **b.** The plasticity phenotypes were calculated for each subcluster and ordered using the same dendrogram as described in Fig. 8. **c**. For comparison the plasticity phenotypes were calculated using previously published data[23] for normal development (2 – 32 wks) **d.** and animals MDed from eye open until either 4, 5, 6, 9 or 32 wks.

Inspection of the plasticity phenotypes identified some obvious and other subtler differences among the subclusters (Figure 9b). Indeed, the pattern of red and green bands in the BD phenotypes was different from 5wk normals (Figure 9) and showed the shift to more GABA_A_α1 and less GluN2A. For the RO subcluster, the light grey bands and number of green bands identified loss of protein expression and a shift to more GluN2A than 2B and more GluA2 than 5wk normals. The RO pattern, however, appeared similar to an older (e.g. 12wk) normally reared animal suggesting that RO may accelerate maturation of these proteins. Thus, these BD and RO treatments led to distinct plasticity phenotypes.

The pattern of red and green bands in the plasticity phenotype for LT-BV and some of the ST-BV subclusters (ST-BV1, ST-BV5) appeared similar to the 5wk normals (Figure 9b) but many of the features were still significantly different from the age-matched normals (Figure 10a, Supplemental Table 10-1). Nonetheless, these subclusters had some consistent differences with less GABA_A_Rs and more GluN2B than 5wk normals. Interestingly, one of the novel features found by PCA, the GluN2A:GluA2 balance, was the only measure where all of the LT-BV subclusters were not different from 5wk normals but both RO and BD were different. Thus, this visualization of the plasticity phenotypes illustrated that the pattern promoted by BV, and LT-BV in particular, was most similar to the normal CP phenotype.

**Figure 10.**
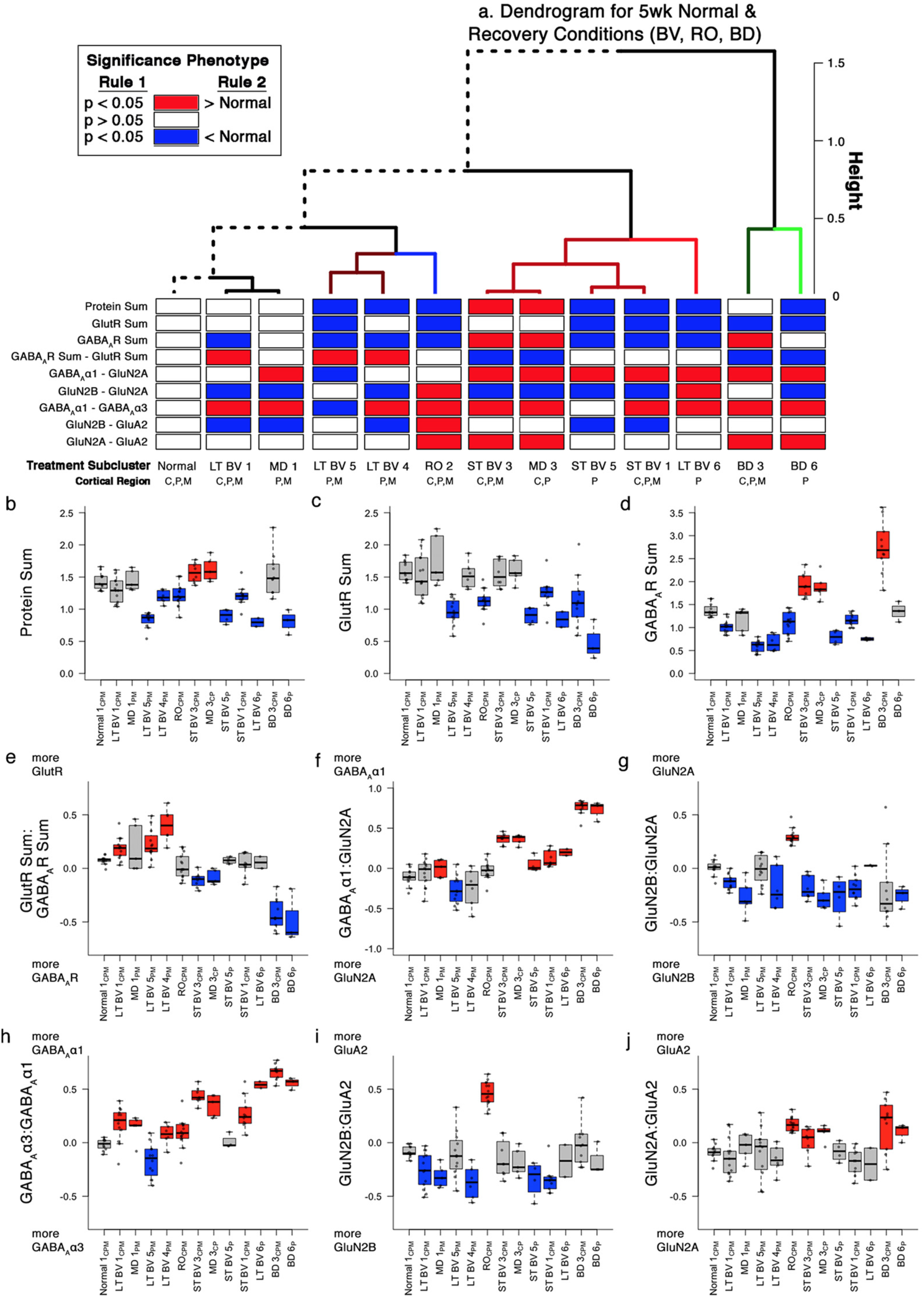
Significant plasticity features. **a**. We used bootstrap analysis to identify plasticity features that were significantly different from 5wk normal animals and color-coded the horizontal bars red if the feature was >normal and blue if it was <normal (p<0.05). b-j. The boxplots show the average protein sum (b-d) and an average index value (e-j) for each of the subclusters. Boxes were colored red if significantly greater than 5wk normals, blue if significantly less than 5wk normals and grey if not significantly different from 5wk normals. For exact Bonferroni corrected p-values see Supplemental Table 10-1.

### Modeling NMDAR and GABA_A_R population kinetics

The subunit composition of NMDARs and GABA_A_Rs help to regulate the threshold for experience-dependent plasticity, in part by controlling receptor kinetics[44], [45]. We used information about receptor kinetics with different subunit compositions to make a model that predicts the average population kinetics and applied it to normal development and the rearing conditions studied here. First, we transformed the 2A:2B and α1:α3 balances into predicted population kinetics (see Methods) and plotted the normal postnatal development (Figure 11 a,b). Both the NMDA and GABA_A_ kinetics rapidly speed up between 2 to 6 weeks of age. Next, we compared the predicted kinetics among the rearing conditions (Figure 11 c, d). The pattern of results is necessarily similar to the balances presented for the indices (Figure 6); however, the predicted kinetics suggests a compression of differences between conditions when the balances favor the mature subunits (2A or α1) versus an accentuation of differences with much slower kinetics when the immature subunits (2B or α3) dominated.

**Figure 11.**
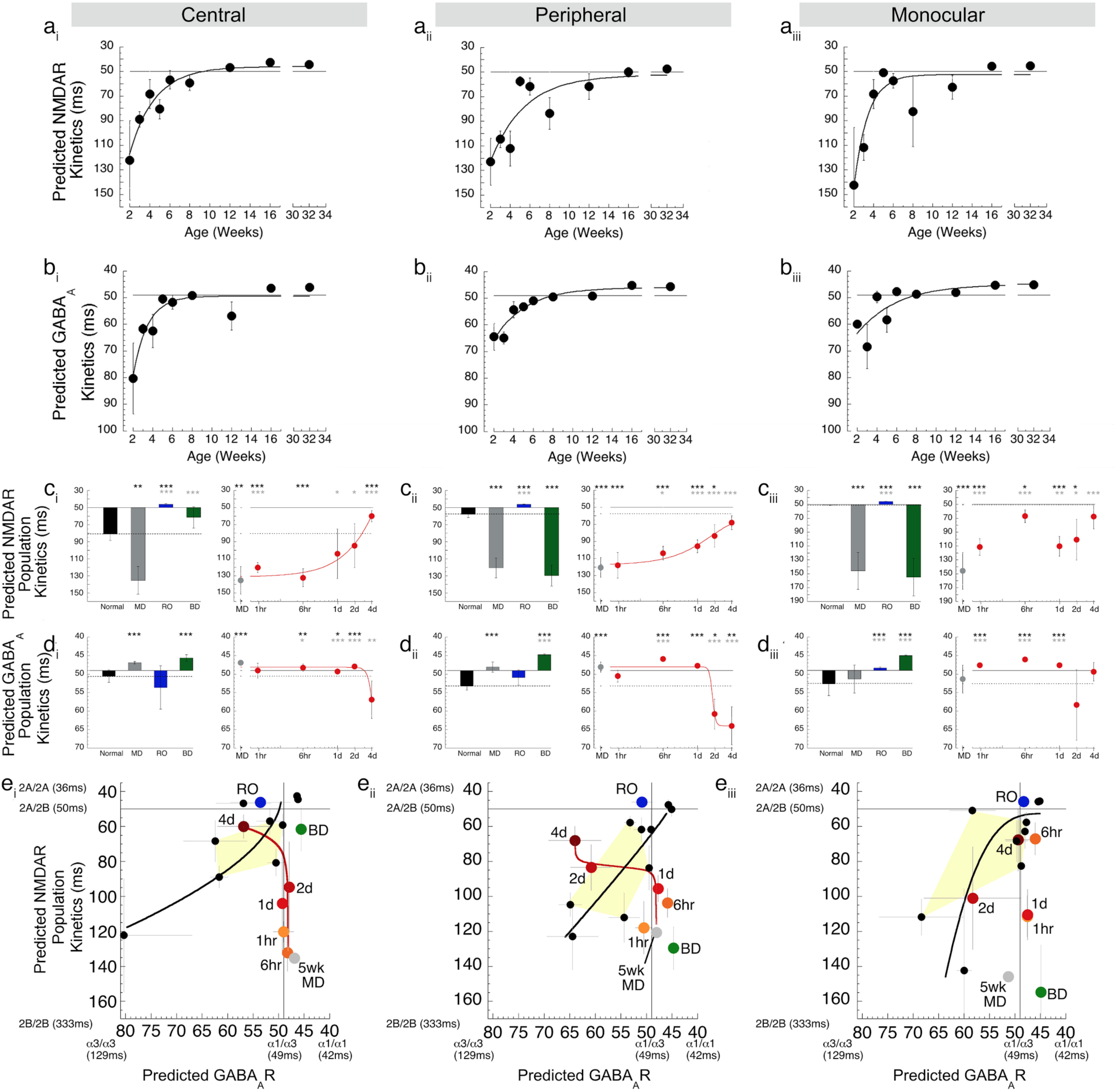
Indices for pairs of receptor subunits and modeling of predicted decay kinetics for a population of NMDARs and GABA_A_Rs. Scatterplots showing the average expression of the predicted population kinetics (**a:**NMDAR, **b:**GABA_A_R) for the regions of V1 (columns) across normal development. Histograms and scatterplots showing the average expression of the predicted population kinetics (**c:**NMDAR, **d:**GABA_A_R) for the regions of V1 (columns) across treatment conditions. **e.** The predicted population kinetics are plotted for both GABA_A_Rs (x-values) and NMDARs (y-values) for normally reared animals age range 2wks-adult with the curve representing the trajectory of the relationship between these features (black dots & line, see **a** and **b**). Also, the data are plotted for 5wk MD (grey dot), RO (blue dot), and BD (green dot). The relationship between NMDAR and GABA_A_R kinetics during BV treatment for 1hr (orange) to 4d (red) is plotted, and the line uses the functions fit to the data in c and d. For exact p-values, Pearson’s R, and equations for the curve-fits see Supplemental Table 11-1.

To address how treatment-induced changes to NMDAR and GABA_A_R composition might affect the relationship between glutamatergic and GABAergic transmission timing we made XY scatterplots using the predicted kinetics (Figure 11 e). During normal development (black line) both balances progressed from slow kinetics at 2wks to faster kinetics through the peak of the CP (Figure 11e-yellow zone, 4-6wks) to reach adult levels. The NMDAR:GABA_A_R kinetics for MD, RO and BD fell outside the window predicted for the normal CP, but in different directions. MD had slower NMDAR (C: 135ms±16ms; P: 121ms±12ms; M: 146ms±27ms) but faster GABA_A_R kinetics (C: 47ms±0.3ms; P: 48ms±1ms; M: 51ms±4ms), RO had faster NMDAR (C: 46ms±0.8ms; P: 46ms±0.4ms; M: 46ms±0.2ms) but normal CP range for GABA_A_R (C:54ms±6ms; P:51ms±2ms; M: 48ms±0.2ms), and BD had faster GABA_A_R (C: 46ms±0.9ms; P: 44ms±0.2ms; M:45ms±0.2ms) but normal CP range NMDAR kinetics in the central region only (C: 61ms±12ms; P: 130ms±12ms; M: 155ms±27ms).

The introduction of BV caused a progressive change in the predicted NMDAR:GABA_A_R kinetics suggesting an initial speeding up of the NMDAR kinetics over the first 1d to 2d followed by a slowing of the GABA_A_R kinetics, especially in the central region. Taken together, the predicted NMDAR:GABA_A_R kinetics provided additional evidence that BV shifts protein expression towards a normal CP balance but none of the treatments reinstated normal kinetics.

## Discussion

Here we studied a set of glutamatergic and GABAergic receptor subunits in V1 that regulate plasticity and explored classifying treatments that cause either persistent bilateral amblyopia (RO or BD) or good acuity in both eyes (BV). Not surprisingly, there was a complex pattern of changes that varied by treatment and region within V1. Applying a new analysis approach, however, using PCA and cluster analysis, identified higher dimensional features and subclusters with different plasticity phenotypes for treatments that promote good versus poor recovery of acuity. The LT-BV plasticity phenotypes were closest to the normal CP pattern while the RO phenotype appeared more similar to an older pattern dominated by GluA2. In contrast, the BD phenotypes were dominated by GABA_A_α1 making it distinct from RO and illustrating that multiple plasticity phenotypes can underlie persistent bilateral amblyopia. The PCA analysis identified an understudied feature, the balance between mature glutamate receptor subunits (GluN2A:GluA2 balance), as a marker that might differentiate treatments supporting good acuity (BV), from those that lead to persistent bilateral amblyopia (RO, BD). Finally, modeling kinetics for NMDAR and GABA_A_R provided additional evidence that BV can return CP-like balances, especially in the central region of V1.

### Study limitations and design

The exploratory nature of the design used here was limited because the small number of animals used leaves unanswered how much variation there is in response to the treatments. The visual manipulations (MD, RO, BD, BV), however, are known to cause consistent changes in visual perception [7], [8], [49]-[52], physiology [7], [29], [31], [53], and molecular mechanisms [23], [27], [54]-[59] that have been reliably measured in a number of laboratories using the cat to study visual system plasticity. Thus, these treatment-induced changes provide an understanding about the pattern of recovery that will be useful for formulating new hypotheses about the links between these proteins and persistent amblyopia.

The study design had some strengths including that (i) the animal model has excellent spatial vision, with a central visual field, so we could compare changes in regions of V1 that represent different parts of the visual field[27], (ii) the treatments were initiated and completed within the CP[34], (iii) there is detailed information about the recovery of physiology for RO and BV[7], [29], [53], [60] and acuity for all 3 treatments[7], [8], [27], [29], [30], (iv) both RO and BD cause persistent bilateral amblyopia[8], [30], and (v) the treatments engage different forms of experience-dependent plasticity (RO: competitive; BD: cooperative with degraded visual input; BV: cooperative with normal visual input).

We observed that only one feature (GluN2A:GluA2 balance) returned to normal after LT-BV treatment raising the hypothesis that it is necessary for good recovery. We were not able to test that question because the molecular tools are not available for manipulating proteins in cat cortex so it will be important to replicate that finding in the mouse and then test the question by directly manipulating those proteins. In addition, a large number of other treatments have been tested to improve recovery after MD, including a brief period of dark-rearing[30], [61], fluoxetine administration[62], perceptual learning[27], [63], or targeting specific molecular mechanisms (e.g., perineuronal nets[64]). Undoubtedly, the timing, length, and type of treatment influences recovery but the conditions used here were necessarily limited because of the labor-intensive nature of this study. Notwithstanding these limitations, the plasticity phenotypes identified RO and BD as different from each other and from normals, but the LT-BV subclusters were remarkably similar to the 5wk normal pattern.

Finally, the design took advantage of the reliability and multiplexing capabilities of Western blotting to obtain a large dataset of plasticity proteins from multiple V1 regions and rearing conditions. Western blotting, however, does not provide information about the cell types, layers, cortical columns, or subcellular localization of these proteins that would reveal which circuits are involved in recovery or persistent amblyopia. Even without that information, the application of high dimensional analyses led to the characterization of features and treatment-based clusters with unique plasticity phenotypes. The phenotyping approach developed here is scalable for studying more proteins or genes, cortical areas, and treatment conditions. Taken together, we think that this approach can be used in other animal models where molecular tools can be combined with visual testing to identify the features and phenotypes necessary for optimal visual recovery.

### BV promoted recovery of CP-like plasticity phenotype and identified GluA2:GluN2A as a balance that differentiated BV treatment

We explored BV treatment because it promotes long-lasting recovery of good acuity in both eyes[27] and those findings are similar to promising results of binocular therapies for treating amblyopia in children[65]. Furthermore, there is good physiological recovery with BV[29], [60]. Thus, it was not surprising to find that LT-BV subclusters had the strongest correlations with normals, or that those subclusters had CP-like phenotypes. However, only one of the features, the GluA2:GluN2A balance, returned to normal levels. Those findings suggest that it may not be necessary to recapitulate every detail of the normal phenotype to support good visual recovery and that the GluA2:GluN2A balance may be a characteristic feature for tracking functional recovery. Although that balance is not commonly quantified, both proteins are critical components of mechanisms regulating experience-dependent plasticity and that balance might signify the adaptive engagement of multiple plasticity mechanisms. For example, the delayed increase in visual responses during ocular dominance plasticity is part of a homeostatic plasticity mechanism regulated by trafficking GluA2-containing AMPARs into the synapse[66], [67]. Meanwhile, the initiation of ocular dominance plasticity requires GluN2A expression[22], and when GluN2A is deleted or reduced MD does not depress deprived eye responses but instead causes enhancement of activity driven by the open eye[21]. Our finding that LT-BV returned a CP-like GluA2:GluN2A balance suggests that BV may prime GluN2A-dependent Hebbian plasticity to consolidate deprived-eye connections while GluA2-dependent homeostatic plasticity enhances deprived-eye responsiveness without triggering runaway excitation[68]-[72]. Thus, the GluA2:GluN2A balance may reflect the idea that during BV treatment the non-deprived eye acts as a *teacher* guiding both cooperative and competitive plasticity mechanisms[29].

### RO versus BD plasticity phenotypes

Because RO and BD treatments cause persistent bilateral amblyopia[7], [8], [30] we expected these conditions to have abnormal phenotypes. We were surprised, however, to find very different phenotypes for these conditions, showing that more than one plasticity phenotype can account for persistent acuity deficits.

RO samples were in a single cluster dominated by an overabundance of GluA2 and more GluN2A than 2B. Together those changes made the RO phenotype appear more similar to an adult than the CP pattern. The increase in GluA2 was in sharp contrast to the loss after BV treatment, and suggests that RO may scale up AMPAR-dependent homeostatic mechanisms to drive recovery[25] without engaging NMDAR-dependent mechanisms to consolidate those changes[73]. Since AMPAR-mediated homeostasis promotes rapid but transient gains in responsiveness [25], [66], [74]-[77] this might explain the labile acuity recovered with RO[7], [8]. Interestingly, the over-representation of GluA2 promoted by RO implicates the dense expression of GluA2-containing synapses at feedback connections onto parvalbumin-positive (PV+) neurons[78]. The feedforward connections onto PV+ neurons may also be involved in RO circuit abnormalities because the labile acuity and early shift to GluN2A after RO are similar to changes found in MeCP2 KOs where an abnormally early shift to GluN2A at synapses onto PV+ neurons that halts acuity development[79], [80]. Taken together, these findings provide preliminary evidence that RO may leave behind feedforward (GluN2A subunits) and feedback abnormalities (GluA2) in PV+ neuron circuits in V1.

Although various models of neural plasticity predict that decreasing firing rates will enhance plasticity that idea has not translated to using BD treatment to improve recovery from MD[30]. BD for weeks or months during the CP has a range of effects on V1 including enhancing the appearance of cytochrome oxidase blobs[81], weakening stimulus evoked activity of PV+ neurons[82] and delaying the developmental increase in GAD65 expression [83]. Here we found that a few days of BD treatment caused an abnormal increase in the expression of GABA_A_α1 throughout V1 and a shift to more GluN2A in the central region. GABA_A_α1 receptors are found on pyramidal cell bodies where PV+ neurons synapse and they serve as regulators of ocular dominance plasticity[20] and the window for coincident spike-time dependent plasticity[24]. A recent study has shown that the loss of PV+ activity caused by BD depends on GABA_A_α1 mechanisms and that blocking this subunit increases BD-evoked activity allowing for LTP of PV+ neurons[84]. Our observation of increased GABA_A_α1 expression suggests that BD treatment may further reduce visually evoked activity in V1 that is compounded by the shift to more GluN2A reducing the availability of NMDA-dependent mechanism needed to consolidate visual recovery.

### Modeling recovery of NMDAR and GABA_A_R kinetics

Our modeling of population kinetics suggests that different physiological changes accompany the 3 treatments. During normal development the increases in NMDAR and GABA_A_R kinetics progress in concert. Physiological studies [85] and our modeling show that this fine balance is decoupled by MD because the delayed shift to GluN2A has slower NMDAR kinetics but the premature increase of GABA_A_α1 has faster GABA_A_R kinetics. Neither RO or BD treatment corrected that decoupling and the modeling suggests that those treatments accelerate the shift to faster adult-like kinetics for NMDARs after RO or GABA_A_Rs after BD. Modeling the kinetics for BV treatment identified 2 phases of recovery especially in the binocular regions of V1. First, between 0-2 days of BV there was a rapid increase in the predicted NMDAR kinetics that was similar to changes that occur between 2 to 4 weeks of age in normal cats. Second, between 2-4 days of BV there was a slowing of the predicted GABA_A_R kinetics and that was opposite to the normal developmental increase in kinetics. These sequential phases of BV treatment do not recapitulate normal development. These results raise the question of whether the BV-driven increase in NMDAR kinetics needs to reach a certain level before triggering the slowing of GABA_A_R kinetics to rebalance these mechanisms. This modeling, however, was based on population data about the expression of the receptor subunits and cannot be extrapolated to individual receptors. Nonetheless, the rapid changes with BV treatment suggest that some aspects of normal development may be missed and it will be important to determine what those are.

### How might these plasticity phenotypes be used for developing and testing treatments for persistent amblyopia?

The distinct plasticity phenotypes classified for RO and BD treatments provide preliminary evidence that multiple neural changes can account for persistent amblyopia and highlight the need to know which mechanisms to target when trying to engage neuroplasticity mechanisms to improve acuity. Whether the treatment should focus on AMPARs, NMDARs, GABA_A_Rs or some combination of those receptors will depend on the underlying plasticity phenotype. Insights into those questions can be addressed in animal models using modern molecular tools and vision testing but translating those findings into treatments for humans will depend on non-invasive ways to determine an individual’s plasticity phenotype. For example, magnetic resonance spectroscopy has been used to measure changes in glutamate or GABA concentrations in human V1 after different types of visual experience (e.g. MD[86]) and receptor expression can be quantified by radioligands labeled for SPECT and PET [87]. New molecular imaging techniques hold the promise of even greater detail with the ability to measure the concentration of receptor subunits[88]-[90]. That information may be comparable to protein analysis in animals models and suitable for constructing plasticity phenotypes for human V1 to facilitate the translation of new treatments. Furthermore, behavioral paradigms linked with specific plasticity mechanisms (e.g. stimulus-selective response plasticity[91]) may further aid in characterizing human plasticity phenotypes. Thus, selecting a treatment to prevent or correct persistent amblyopia may benefit from *in vivo* steps to classify an individual’s plasticity phenotype.

### Conclusions

This exploration of glutamatergic and GABAergic receptor subunit changes in V1 after treatment that promotes either good (BV) or poor (RO, BD) recovery of vision provides a better understanding of the complexity of this problem. Of the treatments studied here, only BV provided evidence for recovery of a CP-like plasticity phenotype in V1. However, only one feature, the GluA2:GluN2A balance, returned to normal levels after BV and that balance is noteworthy because the proteins are regulators of homeostatic and Hebbian plasticity, respectively. The modeling of NMDAR and GABA_A_R kinetics suggests two stages for BV recovery: a rapid increase in NMDAR kinetics, followed by slowing of the predicted GABA_A_R kinetics which together move that balance into the CP range. We identified features of the plasticity phenotypes that may guide future studies on persistent amblyopia to look for high levels of GluA2 and GluN2A following RO, and high levels of GABA_A_α1 after BD treatment. Finally, the plasticity phenotyping is a good approach for uncovering novel neurobiological features that may be important for recovery of acuity and new treatment targets.

## Supplemental Data: Figure Legends

**Figure 7-1 Within group sum of squares with variable cluster sizes.** Scatterplot of the within groups sum of squares was measured across a range of clusters between 2 and 15. An exponential decay fit was applied to the data, and 4τ was taken as the point at which changes in cluster number had little effect on the within groups sum of squares. The optimal number of clusters was identified as 6 (k=6). This value was used to assign the k-means clusters on tSNE reduced data (Figure 7a).

## Supplemental Data: Table Legends

**Table 2-1. The number of Western blot measurements for each cortical region in Normal animals.** Rows summarize the number of runs from the Central (C), Peripheral (P), and Monocular (M) regions of V1 within each age of animal studied. The columns list each of the 7 proteins analyzed using Western blotting. Column sums detail the number of runs across ages and cortical areas.

**Table 2-2. The number of Western blot measurements for each cortical region in MD animals.** Rows summarize the number of runs from the Central (C), Peripheral (P), and Monocular (M) regions of V1 within each age of animal studied. The columns list each of the 7 proteins analyzed using Western blotting. Column sums detail the number of runs across ages and cortical areas.

**Table 3-1. Pearson’s R values in each treatment condition comparing the strength of association between each protein.** The correlation between each protein within a treatment condition was measured (left), and the observed R values are presented in a matrix. This matrix was reordered in Figure 3 to position high R values nearest one another. The Bonferroni corrected p-values (right) were used to identify the most significant correlations between proteins. P-values <0.05 are colored red to simplify identification of significant correlations.

**Table 4-1. Pearson’s R correlations between newly identified plasticity features and PCA dimensions.** The correlation between the PCA scores across all animals, and the first 3 PCA dimensions are presented. P-values of correlations that were significantly correlated after Bonferroni correction are colored red.

**Table 5-1. Table of p-values comparing the values for protein sums and a newly identified plasticity feature in treatment conditions against 5wk Normal animals or 5wk MD animals.** p-values are presented for each cortical area (columns) and plasticity feature (rows). Cortical areas are broken up into comparisons against normal (left) and MD (right). When a curve fit was applied, the equation, degrees of freedom (df), R^2^ value and exact p-value are listed.

**Table 6-1. p-values comparing the each newly identified plasticity feature in treatment conditions against 5wk Normal animals and 5wk MD animals.** p-values are presented for each cortical area (columns) and plasticity feature (rows). Cortical areas are broken up into comparisons against normal (left) and MD (right). When a curve fit was applied, the equation, degrees of freedom (df), R^2^ value and exact p-value are listed.

**Table 8-1. Pearson’s R values comparing the strength of association between each treatment subcluster.** The correlation between each sub-cluster was measured and the observed R values are presented in a matrix. This matrix was reordered in Figure 8 to position high R values nearest one another.

**Table 8-2. Bonferroni corrected p-values between each treatment subcluster.** The Bonferroni corrected p-values were used to identify the most significant correlations between proteins. P-values less than the Bonferroni corrected level (0.0006) are colored red to simplify identification of significant correlations.

**Table 10-1. p-values for each identified plasticity feature within subclusters compared against the Normal animals from cluster 1.** p-values are presented for the Pearson’s R correlations between each plasticity phenotype and the Normal subcluster. The corresponding significance level is indicated by the text color red if the value was significantly above the normal subcluster, and blue if the value was significantly below, and white if not significantly different.

**Table 11-1. p-values for predicted kinetics among treatment conditions compared against the 5 week Normal and 5 week MD animals.** p-values are presented for each cortical area (columns) and for the predicted kinetics of each receptor type(rows). Cortical areas are broken up into comparisons against normal (left) and MD (right). When a curve fit was applied, the equation, degrees of freedom (df), R^2^ value and exact p-value are listed.

## Authors’ contributions

JB Designed research, performed research, analyzed data, wrote/revised the paper; DJ analyzed data, revised the paper; KM designed research, performed research, analyzed data, wrote/revised the paper.

## Acknowledgements

We thank Kyle Hornby and Dr Brett Beston for assistance with data collection.

## Data Availability

The data used to support the findings of this study are available from the corresponding author upon request.

