## Supplemental Materials for "Classification of visual cortex plasticity phenotypes following treatment for amblyopia"

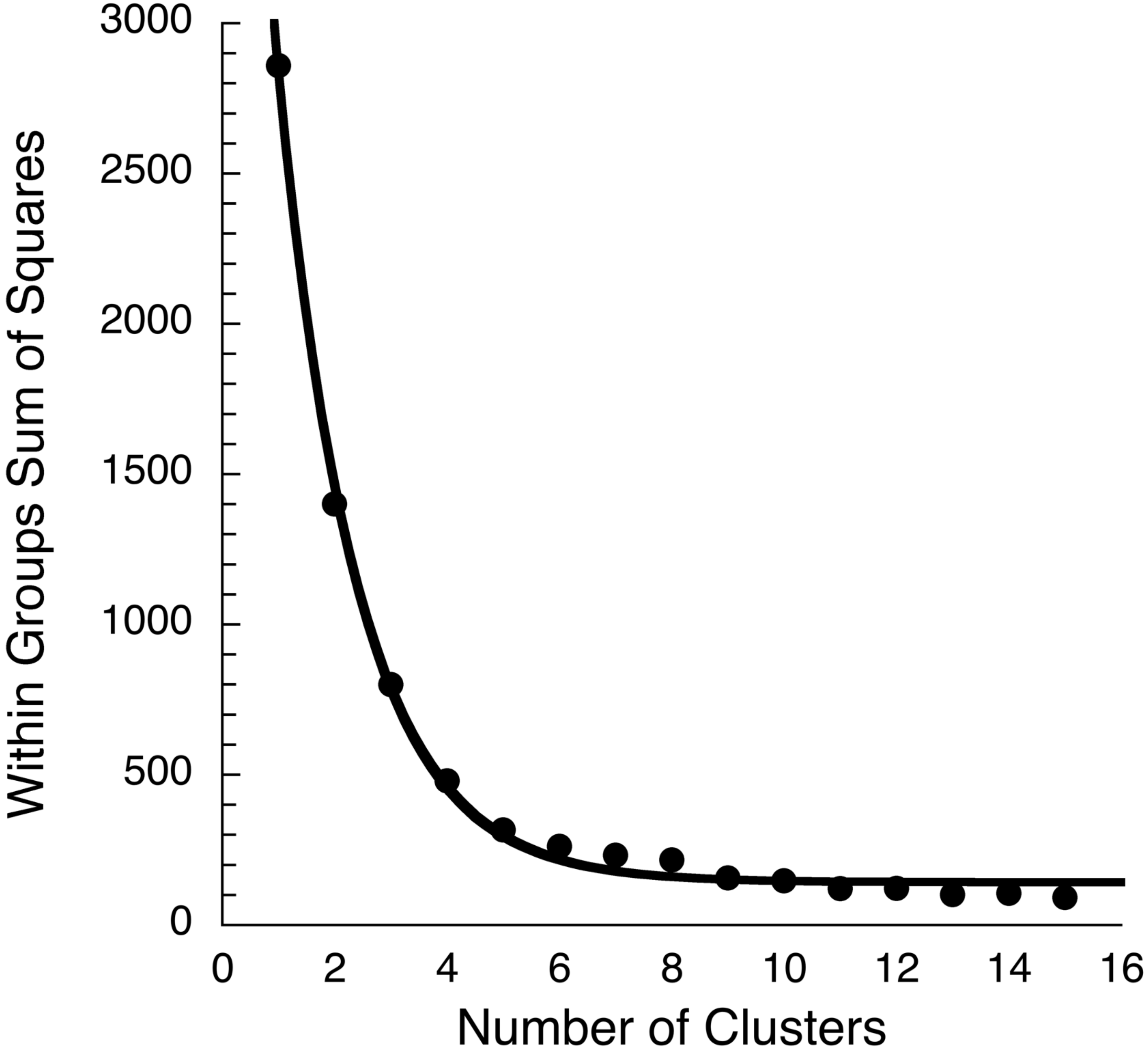


**Figure 7-1 Within group sum of squares with variable cluster sizes.** Scatterplot of the within groups sum of squares was measured across a range of clusters between 2 and 15. An exponential decay fit was applied to the data, and 4τ was taken as the point at which changes in cluster number had little effect on the within groups sum of squares. The optimal number of clusters was identified as 6 (k=6). This value was used to assign the k-means clusters on tSNE reduced data (Figure 7a).

| Weeks BV | Region | GluN1 | GluN2A | GluN2B | $GABA_A\alpha1$ | $GABA_A\alpha3$ | GluA2 | Synapsin |
|---|---|---|---|---|---|---|---|---|
| 2 | C | 4 | 4 | 4 | 4 | 4 | 4 | 4 |
| | P | 16 | 16 | 16 | 16 | 16 | 16 | 16 |
| | M | 4 | 4 | 4 | 4 | 4 | 4 | 4 |
| 3 | C | 4 | 4 | 4 | 4 | 4 | 4 | 4 |
| | P | 16 | 16 | 16 | 16 | 16 | 16 | 16 |
| | M | 4 | 4 | 4 | 4 | 4 | 4 | 4 |
| 4 | C | 4 | 4 | 4 | 4 | 4 | 4 | 4 |
| | P | 16 | 16 | 16 | 16 | 16 | 16 | 16 |
| | M | 4 | 4 | 4 | 4 | 4 | 4 | 4 |
| 5 | C | 4 | 4 | 4 | 4 | 4 | 4 | 4 |
| | P | 16 | 16 | 16 | 15 | 16 | 16 | 16 |
| | M | 4 | 4 | 4 | 4 | 4 | 4 | 4 |
| 6 | C | 4 | 4 | 4 | 4 | 4 | 4 | 4 |
| | P | 16 | 16 | 16 | 16 | 16 | 16 | 16 |
| | M | 4 | 4 | 4 | 4 | 4 | 4 | 4 |
| 8 | C | 6 | 6 | 6 | 6 | 6 | 6 | 6 |
| | P | 22 | 19 | 22 | 22 | 22 | 22 | 22 |
| | M | 2 | 2 | 2 | 2 | 2 | 2 | 2 |
| 12 | C | 4 | 4 | 4 | 4 | 4 | 4 | 4 |
| | P | 16 | 16 | 16 | 16 | 16 | 13 | 16 |
| | M | 4 | 4 | 4 | 4 | 4 | 4 | 4 |
| 16 | C | 4 | 4 | 4 | 4 | 4 | 4 | 4 |
| | P | 16 | 16 | 16 | 16 | 16 | 16 | 16 |
| | M | 4 | 4 | 4 | 4 | 4 | 4 | 4 |
| 32 | C | 4 | 4 | 4 | 4 | 4 | 4 | 4 |
| | P | 16 | 16 | 16 | 16 | 16 | 16 | 16 |
| | M | 4 | 4 | 4 | 4 | 4 | 4 | 4 |
| SUM | | 222 | 219 | 222 | 221 | 222 | 219 | 222 |

**Table 2-1. The number of Western blot measurements for each cortical region in Normal animals.** Rows summarize the number of runs from the Central (C), Peripheral (P), and Monocular (M) regions of V1 within each age of animal studied. The columns list each of the 7 proteins analyzed using Western blotting. Column sums detail the number of runs across ages and cortical areas.

| Weeks MD | Region | GluN1 | GluN2A | GluN2B | GABA$_A$α1 | GABA$_A$α3 | GluA2 | Synapsin |
|---|---|---|---|---|---|---|---|---|
| 4 | C | 4 | 4 | 4 | 4 | 4 | 4 | 4 |
| | P | 16 | 16 | 16 | 16 | 16 | 11 | 16 |
| | M | 4 | 4 | 4 | 4 | 4 | 2 | 4 |
| 5 | C | 6 | 6 | 6 | 6 | 6 | 6 | 4 |
| | P | 18 | 18 | 18 | 18 | 18 | 18 | 11 |
| | M | 4 | 4 | 4 | 4 | 4 | 4 | 4 |
| 6 | C | 8 | 8 | 8 | 8 | 8 | 8 | 8 |
| | P | 32 | 32 | 32 | 32 | 32 | 32 | 32 |
| | M | 6 | 6 | 6 | 6 | 6 | 6 | 6 |
| 9 | C | 8 | 8 | 8 | 8 | 8 | 8 | 8 |
| | P | 36 | 36 | 34 | 34 | 36 | 36 | 31 |
| | M | 8 | 8 | 8 | 6 | 8 | 8 | 6 |
| 32 | C | 4 | 4 | 4 | 4 | 4 | 4 | 4 |
| | P | 16 | 16 | 16 | 16 | 16 | 16 | 16 |
| | M | 2 | 2 | 2 | 2 | 2 | 2 | 2 |
| SUM | | 172 | 172 | 170 | 168 | 172 | 165 | 156 |

**Table 2-2. The number of Western blot measurements for each cortical region in MD animals.** Rows summarize the number of runs from the Central (C), Peripheral (P), and Monocular (M) regions of V1 within each age of animal studied. The columns list each of the 7 proteins analyzed using Western blotting. Column sums detail the number of runs across ages and cortical areas.

| | | Pearson's R | | | | | | | Bonferroni Corrected p-value | | | | | | |
|---|---|---|---|---|---|---|---|---|---|---|---|---|---|---|---|
| | | GluN1 | GluN2A | GluN2B | GABA$_A$α1 | GABA$_A$α3 | GluA2 | Synapsin | GluN1 | GluN2A | GluN2B | GABA$_A$α1 | GABA$_A$α3 | GluA2 | Synapsin |
| 5wk Normal | **GluN1** | 1.0000 | 0.1244 | 0.7605 | **0.8623** | 0.6852 | 0.6861 | **0.8459** | | 14.7036 | 0.0857 | 0.0065 | 0.2927 | 0.2887 | 0.0110 |
| | **GluN2A** | 0.1244 | 1.0000 | 0.4563 | 0.0831 | 0.2667 | 0.4129 | 0.1248 | 14.7036 | | 2.8551 | 16.7446 | 8.4450 | 3.8260 | 14.6803 |
| | **GluN2B** | 0.7605 | 0.4563 | 1.0000 | 0.7056 | 0.6757 | 0.7619 | 0.6297 | 0.0857 | 2.8551 | | 0.2176 | 0.3334 | 0.0836 | 0.5924 |
| | **GABA$_A$α1** | **0.8623** | 0.0831 | 0.7056 | 1.0000 | 0.7158 | 0.6524 | 0.6873 | 0.0065 | 16.7446 | 0.2176 | | 0.1857 | 0.4512 | 0.2839 |
| | **GABA$_A$α3** | 0.6852 | 0.2667 | 0.6757 | 0.7158 | 1.0000 | **0.9535** | 0.7488 | 0.2927 | 8.4450 | 0.3334 | 0.1857 | | 0.0000 | 0.1065 |
| | **GluA2** | 0.6861 | 0.4129 | 0.7619 | 0.6524 | **0.9535** | 1.0000 | 0.7454 | 0.2887 | 3.8260 | 0.0836 | 0.4512 | 0.0000 | | 0.1133 |
| | **Synapsin** | **0.8459** | 0.1248 | 0.6297 | 0.6873 | 0.7488 | 0.7454 | 1.0000 | 0.0110 | 14.6803 | 0.5924 | 0.2839 | 0.1065 | 0.1133 | |
| 5wk MD | **GluN1** | 1.0000 | **0.4100** | **0.6353** | -0.7834 | -0.5047 | 0.0634 | -0.7529 | | 0.0018 | 0.0002 | 3.1854 | 4.6850 | 0.0736 | 16.8673 |
| | **GluN2A** | **0.4100** | 1.0000 | **0.2805** | 0.1070 | -0.0803 | **0.4764** | -0.4470 | 0.0018 | | 0.0006 | 15.3799 | 0.0675 | 0.0000 | 5.4053 |
| | **GluN2B** | **0.6353** | **0.2805** | 1.0000 | -0.3743 | -0.5441 | **0.2998** | -0.2972 | 0.0002 | 0.0006 | | 16.0847 | 2.4495 | 0.0017 | 8.5682 |
| | **GABA$_A$α1** | -0.7834 | 0.1070 | -0.3743 | 1.0000 | 0.5027 | 0.4173 | 0.6579 | 3.1854 | 15.3799 | 16.0847 | | 4.9244 | 9.0319 | 0.1331 |
| | **GABA$_A$α3** | -0.5047 | -0.0803 | -0.5441 | 0.5027 | 1.0000 | -0.0448 | 0.2924 | 4.6850 | 0.0675 | 2.4495 | 4.9244 | | 0.0689 | 4.0109 |
| | **GluA2** | 0.0634 | **0.4764** | **0.2998** | 0.4173 | -0.0448 | 1.0000 | 0.2912 | 0.0736 | 0.0000 | 0.0017 | 9.0319 | 0.0689 | | 0.6893 |
| | **Synapsin** | -0.7529 | -0.4470 | -0.2972 | 0.6579 | 0.2924 | 0.2912 | 1.0000 | 16.8673 | 5.4053 | 8.5682 | 0.1331 | 4.0109 | 0.6893 | |
| RO | **GluN1** | 1.0000 | 0.6894 | 0.4321 | 0.3385 | 0.0863 | 0.1573 | 0.6450 | | 0.1339 | 2.5805 | 4.9665 | 16.1523 | 12.4138 | 0.2679 |
| | **GluN2A** | 0.6894 | 1.0000 | 0.6420 | -0.0955 | -0.0075 | 0.4625 | **0.7514** | 0.1339 | | 0.2794 | 15.6551 | 20.5761 | 2.0130 | 0.0408 |
| | **GluN2B** | 0.4321 | 0.6420 | 1.0000 | 0.1985 | 0.3214 | 0.0836 | 0.3557 | 2.5805 | 0.2794 | | 10.4227 | 5.5118 | 16.3023 | 4.4506 |
| | **GABA$_A$α1** | 0.3385 | -0.0955 | 0.1985 | 1.0000 | 0.5621 | -0.0245 | 0.3207 | 4.9665 | 15.6551 | 10.4227 | | 0.7652 | 19.6111 | 5.5355 |
| | **GABA$_A$α3** | 0.0863 | -0.0075 | 0.3214 | 0.5621 | 1.0000 | -0.0713 | 0.2312 | 16.1523 | 20.5761 | 5.5118 | 0.7652 | | 16.9808 | 8.9548 |
| | **GluA2** | 0.1573 | 0.4625 | 0.0836 | -0.0245 | -0.0713 | 1.0000 | 0.4624 | 12.4138 | 2.0130 | 16.3023 | 19.6111 | 16.9808 | | 2.0148 |
| | **Synapsin** | 0.6450 | **0.7514** | 0.3557 | 0.3207 | 0.2312 | 0.4624 | 1.0000 | 0.2679 | 0.0408 | 4.4506 | 5.5355 | 8.9548 | 2.0148 | |
| BD | **GluN1** | 1.0000 | **0.7646** | **0.7614** | 0.5617 | 0.5436 | **0.7744** | -0.2661 | | 0.0304 | 0.0327 | 0.7683 | 0.9345 | 0.0241 | 7.5155 |
| | **GluN2A** | **0.7646** | 1.0000 | 0.5647 | 0.2690 | 0.1798 | 0.4188 | -0.0405 | 0.0304 | | 0.7433 | 7.3989 | 11.3082 | 2.8587 | 18.7055 |
| | **GluN2B** | **0.7614** | 0.5647 | 1.0000 | 0.5020 | 0.6689 | **0.8751** | -0.3286 | 0.0327 | 0.7433 | | 1.4143 | 0.1868 | 0.0009 | 5.2770 |
| | **GABA$_A$α1** | 0.5617 | 0.2690 | 0.5020 | 1.0000 | **0.7619** | 0.5675 | -0.4707 | 0.7683 | 7.3989 | 1.4143 | | 0.0324 | 0.7197 | 1.8771 |
| | **GABA$_A$α3** | 0.5436 | 0.1798 | 0.6689 | **0.7619** | 1.0000 | **0.7979** | -0.4945 | 0.9345 | 11.3082 | 0.1868 | 0.0324 | | 0.0131 | 1.5173 |
| | **GluA2** | **0.7744** | 0.4188 | **0.8751** | 0.5675 | **0.7979** | 1.0000 | -0.4630 | 0.0241 | 2.8587 | 0.0009 | 0.7197 | 0.0131 | | 2.0049 |
| | **Synapsin** | -0.2661 | -0.0405 | -0.3286 | -0.4707 | -0.4945 | -0.4630 | 1.0000 | 7.5155 | 18.7055 | 5.2770 | 1.8771 | 1.5173 | 2.0049 | |
| 1hr BV | **GluN1** | 1.0000 | 0.3855 | 0.6080 | 0.2625 | 0.3749 | 0.3297 | 0.3648 | | 4.5343 | 0.7552 | 8.6073 | 4.8274 | 6.2014 | 5.1157 |
| | **GluN2A** | 0.3855 | 1.0000 | 0.5765 | 0.4831 | 0.5649 | 0.5557 | -0.3930 | 4.5343 | | 1.0443 | 2.3441 | 1.1682 | 1.2743 | 4.3313 |
| | **GluN2B** | 0.6080 | 0.5765 | 1.0000 | 0.5026 | 0.6824 | 0.6945 | 0.1850 | 0.7552 | 1.0443 | | 2.0127 | 0.3043 | 0.2562 | 11.8611 |
| | **GABA$_A$α1** | 0.2625 | 0.4831 | 0.5026 | 1.0000 | 0.7756 | 0.6488 | -0.3023 | 8.6073 | 2.3441 | 2.0127 | | 0.0636 | 0.4717 | 7.1321 |
| | **GABA$_A$α3** | 0.3749 | 0.5649 | 0.6824 | 0.7756 | 1.0000 | 0.7779 | -0.2078 | 4.8274 | 1.1682 | 0.3043 | 0.0636 | | 0.0607 | 10.8577 |
| | **GluA2** | 0.3297 | 0.5557 | 0.6945 | 0.6488 | 0.7779 | 1.0000 | -0.2947 | 6.2014 | 1.2743 | 0.2562 | 0.4717 | 0.0607 | | 7.4014 |
| | **Synapsin** | 0.3648 | -0.3930 | 0.1850 | -0.3023 | -0.2078 | -0.2947 | 1.0000 | 5.1157 | 4.3313 | 11.8611 | 7.1321 | 10.8577 | 7.4014 | |
| 6hr BV | **GluN1** | 1.0000 | 0.5988 | 0.3296 | **0.7883** | 0.4455 | 0.7283 | -0.7436 | | 0.8326 | 6.2047 | 0.0487 | 3.0797 | 0.1519 | 0.1170 |
| | **GluN2A** | 0.5988 | 1.0000 | 0.3423 | 0.3607 | 0.2277 | 0.6980 | -0.7610 | 0.8326 | | 5.7978 | 5.2382 | 10.0069 | 0.2436 | 0.0849 |
| | **GluN2B** | 0.3296 | 0.3423 | 1.0000 | -0.0006 | 0.3319 | 0.1969 | -0.0005 | 6.2047 | 5.7978 | | 20.9706 | 6.1304 | 11.3330 | 20.9738 |
| | **GABA$_A$α1** | **0.7883** | 0.3607 | -0.0006 | 1.0000 | 0.4846 | 0.6657 | -0.7018 | 0.0487 | 5.2382 | 20.9706 | | 2.3178 | 0.3806 | 0.2301 |
| | **GABA$_A$α3** | 0.4455 | 0.2277 | 0.3319 | 0.4846 | 1.0000 | 0.4798 | -0.1380 | 3.0797 | 10.0069 | 6.1304 | 2.3178 | | 2.4036 | 14.0483 |
| | **GluA2** | 0.7283 | 0.6980 | 0.1969 | 0.6657 | 0.4798 | 1.0000 | **-0.8150** | 0.1519 | 0.2436 | 11.3330 | 0.3806 | 2.4036 | | 0.0260 |
| | **Synapsin** | -0.7436 | -0.7610 | -0.0005 | -0.7018 | -0.1380 | **-0.8150** | 1.0000 | 0.1170 | 0.0849 | 20.9738 | 0.2301 | 14.0483 | 0.0260 | |
| 1d BV | **GluN1** | 1.0000 | 0.6855 | 0.6145 | 0.2300 | **0.9282** | 0.6501 | -0.4983 | | 0.2911 | 0.7035 | 9.9149 | 0.0003 | 0.4639 | 2.0834 |
| | **GluN2A** | 0.6855 | 1.0000 | 0.0939 | -0.1271 | 0.5680 | 0.2423 | -0.3108 | 0.2911 | | 16.2063 | 14.5689 | 1.1342 | 9.4078 | 6.8364 |
| | **GluN2B** | 0.6145 | 0.0939 | 1.0000 | 0.3267 | 0.4719 | 0.4137 | -0.2796 | 0.7035 | 16.2063 | | 6.2991 | 2.5503 | 3.8076 | 7.9555 |
| | **GABA$_A$α1** | 0.2300 | -0.1271 | 0.3267 | 1.0000 | 0.2823 | 0.6785 | -0.4166 | 9.9149 | 14.5689 | 6.2991 | | 7.8546 | 0.3209 | 3.7368 |
| | **GABA$_A$α3** | **0.9282** | 0.5680 | 0.4719 | 0.2823 | 1.0000 | 0.7387 | -0.4850 | 0.0003 | 1.1342 | 2.5503 | 7.8546 | | 0.1275 | 2.3104 |
| | **GluA2** | 0.6501 | 0.2423 | 0.4137 | 0.6785 | 0.7387 | 1.0000 | -0.2997 | 0.4639 | 9.4078 | 3.8076 | 0.3209 | 0.1275 | | 7.2214 |
| | **Synapsin** | -0.4983 | -0.3108 | -0.2796 | -0.4166 | -0.4850 | -0.2997 | 1.0000 | 2.0834 | 6.8364 | 7.9555 | 3.7368 | 2.3104 | 7.2214 | |
| 2d BV | **GluN1** | 1.0000 | 0.5828 | 0.7563 | 0.4025 | 0.5404 | 0.5221 | -0.3177 | | 0.9818 | 0.0929 | 4.0868 | 1.4634 | 1.7143 | 6.6013 |
| | **GluN2A** | 0.5828 | 1.0000 | 0.5211 | 0.3540 | 0.6509 | 0.1215 | 0.1846 | 0.9818 | | 1.7288 | 5.4374 | 0.4596 | 14.8432 | 11.8825 |
| | **GluN2B** | 0.7563 | 0.5211 | 1.0000 | 0.5761 | 0.4723 | 0.2560 | -0.3347 | 0.0929 | 1.7288 | | 1.0490 | 2.5424 | 8.8604 | 6.0401 |
| | **GABA$_A$α1** | 0.4025 | 0.3540 | 0.5761 | 1.0000 | -0.1227 | -0.1081 | -0.2650 | 4.0868 | 5.4374 | 1.0490 | | 14.7830 | 15.4998 | 8.5092 |
| | **GABA$_A$α3** | 0.5404 | 0.6509 | 0.4723 | -0.1227 | 1.0000 | 0.3566 | 0.0017 | 1.4634 | 0.4596 | 2.5424 | 14.7830 | | 5.3577 | 20.9110 |
| | **GluA2** | 0.5221 | 0.1215 | 0.2560 | -0.1081 | 0.3566 | 1.0000 | -0.5480 | 1.7143 | 14.8432 | 8.8604 | 15.4998 | 5.3577 | | 1.3666 |
| | **Synapsin** | -0.3177 | 0.1846 | -0.3347 | -0.2650 | 0.0017 | -0.5480 | 1.0000 | 6.6013 | 11.8825 | 6.0401 | 8.5092 | 20.9110 | 1.3666 | |
| 4d BV | **GluN1** | 1.0000 | 0.7597 | 0.7097 | -0.2995 | 0.2312 | **0.8860** | 0.2433 | | 0.2266 | 0.4518 | 8.4096 | 10.9292 | 0.0135 | 10.4634 |
| | **GluN2A** | 0.7597 | 1.0000 | **0.9083** | -0.1464 | 0.2666 | 0.7004 | 0.3736 | 0.2266 | | 0.0058 | 14.4161 | 9.5874 | 0.5058 | 6.0387 |
| | **GluN2B** | 0.7097 | **0.9083** | 1.0000 | -0.0472 | 0.2951 | 0.7536 | 0.5582 | 0.4518 | 0.0058 | | 18.8348 | 8.5644 | 0.2484 | 1.9639 |
| | **GABA$_A$α1** | -0.2995 | -0.1464 | -0.0472 | 1.0000 | 0.4077 | -0.1449 | 0.4367 | 8.4096 | 14.4161 | 18.8348 | | 5.0863 | 14.4833 | 4.3467 |
| | **GABA$_A$α3** | 0.2312 | 0.2666 | 0.2951 | 0.4077 | 1.0000 | 0.0029 | 0.5863 | 10.9292 | 9.5874 | 8.5644 | 5.0863 | | 20.8653 | 1.5720 |
| | **GluA2** | **0.8860** | 0.7004 | 0.7536 | -0.1449 | 0.0029 | 1.0000 | 0.3818 | 0.0135 | 0.5058 | 0.2484 | 14.4833 | 20.8653 | | 5.8017 |
| | **Synapsin** | 0.2433 | 0.3736 | 0.5582 | 0.4367 | 0.5863 | 0.3818 | 1.0000 | 10.4634 | 6.0387 | 1.9639 | 4.3467 | 1.5720 | 5.8017 | |

**Table 3-1. Pearson's R values in each treatment condition comparing the strength of association between each protein.** The correlation between each protein within a treatment condition was measured (left), and the observed R values are presented in a matrix. This matrix was reordered in Figure 3 to position high R values nearest one another. The Bonferroni corrected p-values (right) were used to identify the most significant correlations between proteins. P-values <0.05 are colored red to simplify identification of significant correlations.

| | All Protein Sum | GlutR Sum | $GABA_AR$ Sum | GluN2B:GluN2A Index | $GABA_A\alpha3:GABA_A\alpha1$ Index | $GABA_A\alpha1$:GluN2A Index | GluN2B:GluA2 Index | GluN2A:GluA2 Index | GlutR Sum:$GABA_AR$ Sum Index | GluA2:GluN1 Index |
|---|---|---|---|---|---|---|---|---|---|---|
| Dim.1 | 0.0000 | 0.0000 | 0.0000 | 1.0000 | 0.0930 | 0.0000 | 1.0000 | 1.0000 | 1.0000 | 1.0000 |
| Dim.2 | 0.7541 | 0.2225 | 0.0000 | 0.0000 | 0.0000 | 0.0000 | 0.0134 | 0.1211 | 1.0000 | 1.0000 |
| Dim.3 | 1.0000 | 0.0000 | 1.0000 | 0.0000 | 0.0336 | 1.0000 | 0.0000 | 0.0000 | 1.0000 | 1.0000 |

**Table 4-1. Pearson's R correlations between newly identified plasticity features and PCA dimensions.** The correlation between the PCA scores across all animals, and the first 3 PCA dimensions are presented. P-values of correlations that were significantly correlated after Bonferroni correction are coloured red.

| | | Central | | | | | Peripheral | | | | | Monocular | | | | |
|---|---|---|---|---|---|---|---|---|---|---|---|---|---|---|---|---|
| | | Vs 5wk Normal | | vs. 5wk MD | | Curve Fit to BV Data | Vs 5wk Normal | | vs. 5wk MD | | Curve Fit to BV Data | Vs 5wk Normal | | vs. 5wk MD | | Curve Fit to BV Data |
| | | Significance | p-value | Significance | p-value | | Significance | p-value | Significance | p-value | | Significance | p-value | Significance | p-value | |
| Total Protein Sum | MD | n.s. | 0.3039 | | | | n.s. | 0.1149 | | | | *** | 0.0000 | | | |
| | RO | n.s. | 0.0963 | n.s. | 0.1663 | | *** | 0.0000 | *** | 0.0000 | | n.s. | 0.4685 | ** | 0.0089 | |
| | BD | n.s. | 0.1937 | n.s. | 0.2379 | | n.s. | 0.2561 | n.s. | 0.1308 | | ** | 0.0077 | n.s. | 0.4364 | |
| | 1hr BV | *** | 0.0006 | ** | 0.0024 | | *** | 0.0000 | *** | 0.0000 | | *** | 0.0000 | *** | 0.0000 | |
| | 6hr BV | n.s. | 0.0825 | n.s. | 0.1652 | | n.s. | 0.1939 | n.s. | 0.3297 | | ** | 0.0048 | n.s. | 0.3811 | |
| | 1d BV | ** | 0.0062 | * | 0.0151 | | *** | 0.0000 | *** | 0.0000 | | n.s. | 0.0664 | *** | 0.0000 | |
| | 2d BV | n.s. | 0.1635 | n.s. | 0.2675 | | *** | 0.0000 | *** | 0.0000 | | *** | 0.0000 | *** | 0.0000 | |
| | 4d BV | n.s. | 0.0965 | n.s. | 0.1760 | | *** | 0.0000 | *** | 0.0000 | | n.s. | 0.0525 | ** | 0.0014 | |
| GlutR Sum | MD | * | 0.0270 | | | | n.s. | 0.4795 | | | | *** | 0.0000 | | | |
| | RO | *** | 0.0000 | ** | 0.0078 | | *** | 0.0000 | *** | 0.0000 | | * | 0.0277 | *** | 0.0000 | |
| | BD | ** | 0.0078 | * | 0.0443 | | *** | 0.0000 | *** | 0.0000 | | *** | 0.0000 | *** | 0.0000 | |
| | 1hr BV | *** | 0.0000 | *** | 0.0000 | | *** | 0.0000 | *** | 0.0000 | | *** | 0.0000 | *** | 0.0000 | |
| | 6hr BV | n.s. | 0.0805 | n.s. | 0.4991 | y = 126.93-44.17*exp(-x/3.15) | n.s. | 0.1701 | n.s. | 0.1937 | | n.s. | 0.4714 | *** | 0.0000 | y = 55.34+64.82*exp(-x/1.20) |
| | 1d BV | n.s. | 0.1737 | n.s. | 0.4264 | df=19 | *** | 0.0000 | *** | 0.0000 | | *** | 0.0002 | *** | 0.0000 | df=22 |
| | 2d BV | n.s. | 0.2705 | * | 0.0303 | $R^2$=0.475 | *** | 0.0000 | *** | 0.0000 | | *** | 0.0000 | *** | 0.0000 | $R^2$=0.584 |
| | 4d BV | n.s. | 0.0671 | ** | 0.0059 | p= 0.0005 | *** | 0.0003 | ** | 0.0010 | | *** | 0.0000 | *** | 0.0000 | p< 0.0001 |
| $GABA_AR$ Sum | MD | ** | 0.0092 | | | | * | 0.0157 | | | | *** | 0.0000 | | | |
| | RO | ** | 0.0029 | *** | 0.0000 | | ** | 0.0085 | *** | 0.0000 | | n.s. | 0.1604 | n.s. | 0.0923 | |
| | BD | n.s. | 0.1868 | n.s. | 0.2901 | | *** | 0.0000 | ** | 0.0067 | | *** | 0.0000 | *** | 0.0000 | |
| | 1hr BV | n.s. | 0.4751 | n.s. | 0.2013 | | *** | 0.0000 | *** | 0.0000 | | ** | 0.0031 | ** | 0.0025 | |
| | 6hr BV | n.s. | 0.1111 | ** | 0.0073 | y = 45.02+58.77*exp(-x/1.37) | * | 0.0104 | n.s. | 0.4545 | y = 39.14+65.54*exp(-x/1.47) | *** | 0.0000 | *** | 0.0000 | |
| | 1d BV | *** | 0.0000 | *** | 0.0000 | df=19 | *** | 0.0000 | *** | 0.0000 | df=78 | * | 0.0217 | *** | 0.0000 | |
| | 2d BV | *** | 0.0000 | *** | 0.0000 | $R^2$=0.581 | *** | 0.0000 | *** | 0.0000 | $R^2$=0.360 | *** | 0.0000 | * | 0.0104 | |
| | 4d BV | *** | 0.0000 | *** | 0.0000 | p< 0.0001 | *** | 0.0000 | *** | 0.0000 | p< 0.0001 | n.s. | 0.4082 | n.s. | 0.1277 | |
| $GABA_AR$ Sum:GlutR Sum Index | MD | ** | 0.0014 | | | | ** | 0.0088 | | | | *** | 0.0000 | | | |
| | RO | n.s. | 0.3428 | ** | 0.0060 | | ** | 0.0011 | n.s. | 0.2762 | | n.s. | 0.3866 | *** | 0.0000 | |
| | BD | ** | 0.0072 | n.s. | 0.0656 | | *** | 0.0000 | *** | 0.0000 | | *** | 0.0000 | *** | 0.0000 | |
| | 1hr BV | * | 0.0161 | n.s. | 0.3428 | | * | 0.0232 | n.s. | 0.3166 | | n.s. | 0.3530 | *** | 0.0000 | |
| | 6hr BV | n.s. | 0.4307 | * | 0.0274 | y = 0.73-0.76*exp(-x/3.37) | *** | 0.0005 | n.s. | 0.2605 | y = 0.44-0.46*exp(-x/2.68) | *** | 0.0000 | *** | 0.0000 | |
| | 1d BV | *** | 0.0000 | *** | 0.0000 | df=19 | *** | 0.0008 | *** | 0.0000 | df=78 | n.s. | 0.2638 | *** | 0.0000 | |
| | 2d BV | *** | 0.0000 | *** | 0.0000 | $R^2$=0.716 | *** | 0.0005 | *** | 0.0000 | $R^2$=0.49 | n.s. | 0.2473 | *** | 0.0001 | |
| | 4d BV | *** | 0.0002 | *** | 0.0000 | p< 0.0001 | *** | 0.0000 | *** | 0.0000 | p< 0.0001 | * | 0.0177 | *** | 0.0000 | |

**Table 5-1. Table of p-values comparing the values for protein sums and a newly identified plasticity feature in treatment conditions against 5wk Normal animals or 5wk MD animals.** p-values are presented for each cortical area (columns) and plasticity feature (rows). Cortical areas are broken up into comparisons against normal (left) and MD (right). When a curve fit was applied, the equation, degrees of freedom (df), R2 value and exact p-value are listed.

| | | Central | | | | | Peripheral | | | | | Monocular | | | | |
|---|---|---|---|---|---|---|---|---|---|---|---|---|---|---|---|---|
| | | Vs 5wk Normal | | vs. 5wk MD | | Curve Fit to BV Data | Vs 5wk Normal | | vs. 5wk MD | | Curve Fit to BV Data | Vs 5wk Normal | | vs. 5wk MD | | Curve Fit to BV Data |
| | | Significance | p-value | Significance | p-value | | Significance | p-value | Significance | p-value | | Significance | p-value | Significance | p-value | |
| **GABA$_A$α1:GluN2A Index** | **MD** | *** | 0.0000 | | | | *** | 0.0000 | | | | n.s. | 0.2515 | | | |
| | **RO** | n.s. | 0.3777 | *** | 0.0000 | | ** | 0.0017 | *** | 0.0000 | | n.s. | 0.3474 | n.s. | 0.4161 | |
| | **BD** | ** | 0.0038 | n.s. | 0.3288 | | *** | 0.0000 | *** | 0.0000 | | *** | 0.0000 | *** | 0.0000 | |
| | **1hr BV** | * | 0.0183 | * | 0.0280 | | *** | 0.0000 | ** | 0.0046 | | *** | 0.0003 | *** | 0.0002 | |
| | **6hr BV** | ** | 0.0014 | *** | 0.0000 | y = 0.89-1.15*exp(-x/3.86) | *** | 0.0000 | n.s. | 0.1960 | y = 0.36-0.63*exp(-x/1.87) | *** | 0.0000 | *** | 0.0000 | |
| | **1d BV** | * | 0.0197 | *** | 0.0000 | df=22 | ** | 0.0018 | *** | 0.0000 | df=89 | *** | 0.0000 | *** | 0.0000 | |
| | **2d BV** | n.s. | 0.1635 | *** | 0.0000 | $R^2$=0.732 | n.s. | 0.4574 | *** | 0.0000 | $R^2$=0.390 | n.s. | 0.5014 | n.s. | 0.4429 | |
| | **4d BV** | *** | 0.0004 | *** | 0.0000 | p< 0.0001 | * | 0.0484 | *** | 0.0000 | p< 0.0001 | * | 0.0319 | * | 0.0242 | |
| **GluN2B:GluN2A Index** | **MD** | ** | 0.00304 | | | | *** | 0 | | | | *** | 0.00067 | | | |
| | **RO** | *** | 0 | *** | 0 | | *** | 0 | *** | 0 | | *** | 0 | *** | 0 | |
| | **BD** | * | 0.04703 | *** | 0.00011 | | *** | 0.00076 | n.s. | 0.42429 | | *** | 0 | n.s. | 0.33648 | |
| | **1hr BV** | *** | 0 | *** | 0.00078 | | *** | 0 | n.s. | 0.31989 | | *** | 0 | *** | 0.00066 | |
| | **6hr BV** | *** | 0 | n.s. | 0.36396 | y = 0.63-0.92*exp(-x/10.98) | *** | 0 | n.s. | 0.0559 | y = 0.02+-0.26*exp(-x/2.09) | n.s. | 0.12763 | *** | 0 | y = -0.08-0.32*exp(-x/0.04) |
| | **1d BV** | n.s. | 0.12805 | * | 0.03144 | df=23 | *** | 0 | ** | 0.00142 | df=77 | *** | 0.00001 | ** | 0.00294 | df=24 |
| | **2d BV** | n.s. | 0.38994 | * | 0.02734 | $R^2$=0.458 | * | 0.03331 | ** | 0.00679 | $R^2$=0.16 | n.s. | 0.10927 | * | 0.0435 | $R^2$=0.240 |
| | **4d BV** | * | 0.01279 | *** | 0 | p=0.0002 | * | 0.03347 | *** | 0 | p= 0.0003 | n.s. | 0.29108 | *** | 0.00027 | p= 0.0111 |
| **GABA$_A$α3:GABA$_A$α1 Index** | **MD** | *** | 0 | | | | *** | 0 | | | | n.s. | 0.09922 | | | |
| | **RO** | n.s. | 0.30873 | * | 0.02197 | | ** | 0.00832 | *** | 0 | | *** | 0.00081 | n.s. | 0.12593 | |
| | **BD** | *** | 0.00034 | n.s. | 0.05618 | | *** | 0 | *** | 0 | | *** | 0 | *** | 0 | |
| | **1hr BV** | * | 0.02394 | n.s. | 0.08849 | | *** | 0.00036 | *** | 0.00002 | | *** | 0 | *** | 0 | |
| | **6hr BV** | * | 0.02068 | * | 0.03329 | y = -0.29+(0.68+0.29)/(1+(x/0.09)^0.19) | *** | 0 | *** | 0.00008 | y = -0.16+(0.29+0.16)/(1+(x/1.76)^3.01) | *** | 0 | *** | 0 | |
| | **1d BV** | n.s. | 0.06521 | *** | 0 | df=24 | *** | 0 | ** | 0.0022 | df=91 | ** | 0.00456 | n.s. | 0.13167 | |
| | **2d BV** | *** | 0.00001 | *** | 0 | $R^2$=0.29 | n.s. | 0.26614 | ** | 0.00309 | $R^2$=0.250 | n.s. | 0.28274 | * | 0.04251 | |
| | **4d BV** | n.s. | 0.50054 | ** | 0.00386 | p= 0.0045 | n.s. | 0.11633 | *** | 0 | p< 0.0001 | n.s. | 0.07932 | n.s. | 0.21442 | |
| **GluN2B:GluA2 Index** | **MD** | n.s. | 0.1269 | | | | *** | 0.0000 | | | | n.s. | 0.0502 | | | |
| | **RO** | *** | 0.0000 | *** | 0.0000 | | *** | 0.0000 | *** | 0.0000 | | *** | 0.0000 | *** | 0.0000 | |
| | **BD** | n.s. | 0.2035 | ** | 0.0060 | | n.s. | 0.3953 | *** | 0.0001 | | n.s. | 0.2542 | ** | 0.0084 | |
| | **1hr BV** | ** | 0.0041 | n.s. | 0.0503 | | *** | 0.0000 | n.s. | 0.2301 | | ** | 0.0014 | n.s. | 0.0559 | |
| | **6hr BV** | *** | 0.0000 | *** | 0.0000 | y = 1.15-1.52*exp(-(x-1.61)^ 2/-5.99^2 | *** | 0.0006 | * | 0.0282 | y = -0.01-0.34*exp(-(x-0.83)^ 2/2.06^2 | * | 0.0117 | *** | 0.0000 | |
| | **1d BV** | *** | 0.0007 | n.s. | 0.0942 | df=25 | *** | 0.0000 | n.s. | 0.0559 | df=92 | n.s. | 0.1674 | *** | 0.0000 | |
| | **2d BV** | *** | 0.0000 | *** | 0.0000 | $R^2$=0.231 | *** | 0.0005 | n.s. | 0.0710 | $R^2$=0.16 | n.s. | 0.4401 | n.s. | 0.0624 | |
| | **4d BV** | n.s. | 0.3750 | * | 0.0400 | p=0.0112 | n.s. | 0.3682 | *** | 0.0000 | p< 0.0001 | n.s. | 0.0505 | *** | 0.0000 | |
| **GluN2A:GluA2 Index** | **MD** | n.s. | 0.2445 | | | | n.s. | 0.3563 | | | | *** | 0.0000 | na | na | |
| | **RO** | *** | 0.0009 | n.s. | 0.0535 | | *** | 0.0000 | *** | 0.0000 | | *** | 0.0000 | *** | 0.0000 | |
| | **BD** | * | 0.0362 | *** | 0.0004 | | *** | 0.0000 | *** | 0.0000 | | *** | 0.0002 | * | 0.0125 | |
| | **1hr BV** | n.s. | 0.2076 | * | 0.0244 | | n.s. | 0.4150 | n.s. | 0.4175 | | n.s. | 0.0716 | *** | 0.0006 | |
| | **6hr BV** | n.s. | 0.2357 | ** | 0.0027 | | n.s. | 0.0790 | n.s. | 0.1535 | | *** | 0.0000 | * | 0.0294 | |
| | **1d BV** | *** | 0.0000 | *** | 0.0000 | | * | 0.0434 | * | 0.0127 | | ** | 0.0016 | n.s. | 0.1044 | |
| | **2d BV** | *** | 0.0000 | *** | 0.0000 | | n.s. | 0.1986 | n.s. | 0.1083 | | n.s. | 0.0887 | n.s. | 0.4977 | |
| | **4d BV** | n.s. | 0.0616 | *** | 0.0007 | | n.s. | 0.1366 | n.s. | 0.1858 | | n.s. | 0.4026 | * | 0.0308 | |

**Table 6-1. p-values comparing the each newly identified plasticity feature in treatment conditions against 5wk Normal animals and 5wk MD animals.** p-values are presented for each cortical area (columns) and plasticity feature (rows). Cortical areas are broken up into comparisons against normal (left) and MD (right). When a curve fit was applied, the equation, degrees of freedom (df), R2 value and exact p-value are listed.

| | Normal 1 | MD 1 | ST BV 1 | LT BV 1 | RO 2 | MD 3 | ST BV 3 | BD 3 | LT BV 4 | ST BV 5 | LT BV 5 | LT BV 6 | BD 6 |
|---|---|---|---|---|---|---|---|---|---|---|---|---|---|
| **Normal 1** | 1 | 0.97601795 | 0.96105886 | 0.98117 | 0.95768142 | 0.94818068 | 0.94383317 | 0.78413725 | 0.95461708 | 0.96535462 | 0.98297823 | 0.85942578 | 0.62173468 |
| **MD 1** | 0.97601795 | 1 | 0.97744298 | 0.99592531 | 0.88914347 | 0.95365077 | 0.94408703 | 0.76075697 | 0.98352939 | 0.9857111 | 0.95770586 | 0.89630038 | 0.65319854 |
| **ST BV 1** | 0.96105886 | 0.97744298 | 1 | 0.98000413 | 0.8578434 | 0.98158073 | 0.9826135 | 0.85467041 | 0.94144785 | 0.9899475 | 0.9110114 | 0.95290107 | 0.77160096 |
| **LT BV 1** | 0.98117 | 0.99592531 | 0.98000413 | 1 | 0.90193003 | 0.94544405 | 0.94037068 | 0.75248635 | 0.98758197 | 0.97844821 | 0.96541071 | 0.91441834 | 0.63534331 |
| **RO 2** | 0.95768142 | 0.88914347 | 0.8578434 | 0.90193003 | 1 | 0.85699952 | 0.85137272 | 0.69798422 | 0.8761915 | 0.8668415 | 0.9631905 | 0.72680318 | 0.47573671 |
| **MD 3** | 0.94818068 | 0.95365077 | 0.98158073 | 0.94544405 | 0.85699952 | 1 | 0.99775672 | 0.91971517 | 0.88803935 | 0.98380715 | 0.88099307 | 0.90030944 | 0.83242446 |
| **ST BV 3** | 0.94383317 | 0.94408703 | 0.9826135 | 0.94037068 | 0.85137272 | 0.99775672 | 1 | 0.93098098 | 0.87805939 | 0.97593087 | 0.87093562 | 0.91606814 | 0.84324473 |
| **BD 3** | 0.78413725 | 0.76075697 | 0.85467041 | 0.75248635 | 0.69798422 | 0.91971517 | 0.93098098 | 1 | 0.64415169 | 0.84073526 | 0.66177201 | 0.79637235 | 0.94438398 |
| **LT BV 4** | 0.95461708 | 0.98352939 | 0.94144785 | 0.98758197 | 0.8761915 | 0.88803935 | 0.87805939 | 0.64415169 | 1 | 0.94535345 | 0.96449423 | 0.87660664 | 0.52623206 |
| **ST BV 5** | 0.96535462 | 0.9857111 | 0.9899475 | 0.97844821 | 0.8668415 | 0.98380715 | 0.97593087 | 0.84073526 | 0.94535345 | 1 | 0.92400086 | 0.90578365 | 0.75550348 |
| **LT BV 5** | 0.98297823 | 0.95770586 | 0.9110114 | 0.96541071 | 0.9631905 | 0.88099307 | 0.87093562 | 0.66177201 | 0.96449423 | 0.92400086 | 1 | 0.79577094 | 0.47606361 |
| **LT BV 6** | 0.85942578 | 0.89630038 | 0.95290107 | 0.91441834 | 0.72680318 | 0.90030944 | 0.91606814 | 0.79637235 | 0.87660664 | 0.90578365 | 0.79577094 | 1 | 0.76469463 |
| **BD 6** | 0.62173468 | 0.65319854 | 0.77160096 | 0.63534331 | 0.47573671 | 0.83242446 | 0.84324473 | 0.94438398 | 0.52623206 | 0.75550348 | 0.47606361 | 0.76469463 | 1 |

**Table 8-1. Pearson's R values comparing the strength of association between each treatment subcluster.** The correlation between each sub-cluster was measured and the observed R values are presented in a matrix. This matrix was reordered in Figure 8 to position high R values nearest one another.

| | Normal 1 | MD 1 | ST BV 1 | LT BV 1 | RO 2 | MD 3 | ST BV 3 | BD 3 | LT BV 4 | ST BV 5 | LT BV 5 | LT BV 6 | BD 6 |
|---|---|---|---|---|---|---|---|---|---|---|---|---|---|
| **Normal 1** | NA | 0.00003 | 0.00014 | 0.00002 | 0.00018 | 0.00033 | 0.00042 | 0.02125 | 0.00023 | 0.0001 | 0.00001 | 0.00623 | 0.09983 |
| **MD 1** | 0.00003 | NA | 0.00003 | 0 | 0.00313 | 0.00024 | 0.00042 | 0.02839 | 0.00001 | 0.00001 | 0.00018 | 0.00258 | 0.07903 |
| **ST BV 1** | 0.00014 | 0.00003 | NA | 0.00002 | 0.00644 | 0.00002 | 0.00001 | 0.00686 | 0.00048 | 0 | 0.00165 | 0.00025 | 0.02492 |
| **LT BV 1** | 0.00002 | 0 | 0.00002 | NA | 0.00219 | 0.00039 | 0.00051 | 0.03122 | 0 | 0.00002 | 0.0001 | 0.00147 | 0.09049 |
| **RO 2** | 0.00018 | 0.00313 | 0.00644 | 0.00219 | NA | 0.00655 | 0.00732 | 0.05421 | 0.00431 | 0.00533 | 0.00012 | 0.0411 | 0.23344 |
| **MD 3** | 0.00033 | 0.00024 | 0.00002 | 0.00039 | 0.00655 | NA | 0 | 0.00122 | 0.00322 | 0.00001 | 0.00385 | 0.0023 | 0.01034 |
| **ST BV 3** | 0.00042 | 0.00042 | 0.00001 | 0.00051 | 0.00732 | 0 | NA | 0.00078 | 0.00413 | 0.00003 | 0.00487 | 0.00139 | 0.00853 |
| **BD 3** | 0.02125 | 0.02839 | 0.00686 | 0.03122 | 0.05421 | 0.00122 | 0.00078 | NA | 0.08473 | 0.00893 | 0.07385 | 0.01802 | 0.00041 |
| **LT BV 4** | 0.00023 | 0.00001 | 0.00048 | 0 | 0.00431 | 0.00322 | 0.00413 | 0.08473 | NA | 0.00039 | 0.00011 | 0.00427 | 0.18034 |
| **ST BV 5** | 0.0001 | 0.00001 | 0 | 0.00002 | 0.00533 | 0.00001 | 0.00003 | 0.00893 | 0.00039 | NA | 0.00104 | 0.00195 | 0.03017 |
| **LT BV 5** | 0.00001 | 0.00018 | 0.00165 | 0.0001 | 0.00012 | 0.00385 | 0.00487 | 0.07385 | 0.00011 | 0.00104 | NA | 0.01817 | 0.23308 |
| **LT BV 6** | 0.00623 | 0.00258 | 0.00025 | 0.00147 | 0.0411 | 0.0023 | 0.00139 | 0.01802 | 0.00427 | 0.00195 | 0.01817 | NA | 0.02709 |
| **BD 6** | 0.09983 | 0.07903 | 0.02492 | 0.09049 | 0.23344 | 0.01034 | 0.00853 | 0.00041 | 0.18034 | 0.03017 | 0.23308 | 0.02709 | NA |

**Table 8-2. Bonferroni corrected p-values between each treatment subcluster.** The Bonferroni corrected p-values were used to identify the most significant correlations between proteins. P-values less than the Bonferroni corrected level (0.0006) are coloured red to simplify identification of significant correlations.

| | Comparison | MD 1 | LT BV 1 | ST BV 1 | RO 2 | MD 3 | ST BV 3 | BD 3 | LT BV 4 | ST BV 5 | LT BV 5 | LT BV 6 | BD 6 |
|---|---|---|---|---|---|---|---|---|---|---|---|---|---|
| **All Protein Sum** | pvalue | 0 | 0 | 0 | 0 | 0 | 0 | 0 | 0 | 0.01366 | 0.00003 | 0 | 0.00062 |
| | Asterisk | *** | *** | *** | *** | *** | *** | *** | *** | * | *** | *** | *** |
| | color | red | red | red | red | red | red | red | red | blue | blue | blue | blue |
| **GlutR Sum** | pvalue | 0 | 0 | 0 | 0 | 0 | 0 | 0.00006 | 0 | 0.00001 | 0 | 0.03855 | 0.00004 |
| | Asterisk | *** | *** | *** | *** | *** | *** | *** | *** | *** | *** | * | *** |
| | color | red | red | red | red | red | red | red | red | red | red | red | blue |
| **GABA$_A$R Sum** | pvalue | 0.00281 | 0 | 0 | 0 | 0 | 0 | 0 | 0 | 0 | 0 | 0 | 0.39037 |
| | Asterisk | ** | *** | *** | *** | *** | *** | *** | *** | *** | *** | *** | n.s. |
| | color | blue | blue | blue | blue | red | red | red | blue | blue | blue | blue | white |
| **GlutR:GABA$_A$R Sum** | pvalue | 0 | 0 | 0 | 0 | 0 | 0 | 0 | 0 | 0 | 0 | 0 | 0.0001 |
| | Asterisk | *** | *** | *** | *** | *** | *** | *** | *** | *** | *** | *** | *** |
| | color | red | red | red | red | red | red | blue | red | red | red | red | blue |
| **GABA$_A$α1:GluN2A** | pvalue | 0 | 0 | 0 | 0 | 0 | 0 | 0 | 0.27593 | 0 | 0.41887 | 0 | 0 |
| | Asterisk | *** | *** | *** | *** | *** | *** | *** | n.s. | *** | n.s. | *** | *** |
| | color | red | red | red | red | red | red | red | white | red | white | red | red |
| **GluN2B:GluN2A** | pvalue | 0.00206 | 0.02562 | 0.17913 | 0 | 0.00009 | 0.2111 | 0.30192 | 0.31158 | 0.00968 | 0 | 0 | 0.00003 |
| | Asterisk | ** | * | n.s. | *** | *** | n.s. | n.s. | n.s. | ** | *** | *** | *** |
| | color | blue | red | white | red | blue | white | white | white | blue | red | red | blue |
| **GABA$_A$α1:GABA$_A$α3** | pvalue | 0.00017 | 0.00009 | 0 | 0.0205 | 0 | 0 | 0 | 0.03374 | 0.02511 | 0 | 0 | 0 |
| | Asterisk | *** | *** | *** | * | *** | *** | *** | * | * | *** | *** | *** |
| | color | red | red | red | red | red | red | red | red | blue | blue | red | red |
| **GluN2B:GluA2** | pvalue | 0 | 0 | 0.00011 | 0 | 0 | 0 | 0 | 0.01204 | 0.00196 | 0 | 0 | 0 |
| | Asterisk | *** | *** | *** | *** | *** | *** | *** | * | ** | *** | *** | *** |
| | color | red | red | red | red | red | red | red | red | red | red | red | red |
| **GluN2A:GluA2** | pvalue | 0 | 0 | 0 | 0 | 0 | 0 | 0 | 0 | 0 | 0.00001 | 0.01706 | 0 |
| | Asterisk | *** | *** | *** | *** | *** | *** | *** | *** | *** | *** | * | *** |
| | color | red | red | red | red | red | red | red | red | red | red | red | red |

**Table 10-1. p-values for each identified plasticity feature within subclusters compared against the Normal animals from cluster 1.**p-values are presented for the Pearson's R correlations between each plasticity phenotype and the Normal subcluster. The corresponding significance level is indicated by the text colour red if the value was significantly above the normal subcluster, and blue if the value was significantly below, and white if not significantly different.

| | | Central | | | | | Peripheral | | | | | Monocular | | | | |
|---|---|---|---|---|---|---|---|---|---|---|---|---|---|---|---|---|
| | | Vs 5wk Normal | | vs. 5wk MD | | Curve Fit to BV Data | Vs 5wk Normal | | vs. 5wk MD | | Curve Fit to BV Data | Vs 5wk Normal | | vs. 5wk MD | | Curve Fit to BV Data |
| | | Significance | p-value | Significance | p-value | | Significance | p-value | Significance | p-value | | Significance | p-value | Significance | p-value | |
| **NMDAR Predicted Decay Kinetics** | **MD** | ** | 0.00312 | | | | *** | 0 | | | | *** | 0.0007 | | | |
| | **RO** | *** | 0 | *** | 0 | | *** | 0 | *** | 0 | | *** | 0 | *** | 0 | |
| | **BD** | n.s. | 0.0836 | *** | 0 | | *** | 0 | n.s. | 0.22522 | | *** | 0 | n.s. | 0.33676 | |
| | **1hr BV** | *** | 0 | *** | 0.00082 | | *** | 0 | n.s. | 0.42141 | | *** | 0 | *** | 0.00053 | |
| | **6hr BV** | *** | 0 | n.s. | 0.36066 | y = -126.4+257.7*exp(-x/12.41) | *** | 0 | * | 0.01421 | y = 63.9+53.77*exp(-x/1.76) | * | 0.03776 | *** | 0 | |
| | **1d BV** | n.s. | 0.12732 | * | 0.03145 | df=23 | *** | 0 | *** | 0.00002 | df=87 | *** | 0 | ** | 0.00303 | |
| | **2d BV** | n.s. | 0.29523 | * | 0.02565 | $R^2$=0.444 | * | 0.01772 | *** | 0.00063 | $R^2$=0.153 | * | 0.0441 | * | 0.04309 | |
| | **4d BV** | ** | 0.00108 | *** | 0 | p= 0.0003 | n.s. | 0.07203 | *** | 0 | p= 0.0002 | n.s. | 0.17444 | *** | 0 | |
| **$GABA_AR$ Predicted Decay Kinetics** | **MD** | *** | 0 | | | | *** | 0.00035 | | | | n.s. | 0.37952 | | | |
| | **RO** | n.s. | 0.30147 | n.s. | 0.08113 | | n.s. | 0.15855 | n.s. | 0.0877 | | *** | 0 | *** | 0 | |
| | **BD** | *** | 0 | n.s. | 0.05567 | | *** | 0 | *** | 0 | | *** | 0 | *** | 0 | |
| | **1hr BV** | n.s. | 0.22184 | n.s. | 0.09753 | | n.s. | 0.05472 | n.s. | 0.05363 | | *** | 0 | *** | 0 | |
| | **6hr BV** | ** | 0.00871 | * | 0.04659 | y = 448.9+(48.16-448.9)/(1+(x/6.49)^7.85) | *** | 0 | *** | 0 | y = 63.99+(48.05-63.99)/(1+(x/1.84)^16.59) | *** | 0 | *** | 0 | |
| | **1d BV** | * | 0.02853 | *** | 0.00001 | df=24 | *** | 0 | n.s. | 0.09948 | df=91 | *** | 0 | *** | 0 | |
| | **2d BV** | *** | 0 | *** | 0.00001 | $R^2$=0.39 | * | 0.02597 | *** | 0.00013 | $R^2$=0.327 | n.s. | 0.27708 | n.s. | 0.20512 | |
| | **4d BV** | n.s. | 0.10545 | ** | 0.00805 | p=0.0006 | ** | 0.00938 | *** | 0.00003 | p< 0.0001 | n.s. | 0.10039 | n.s. | 0.19565 | |

**Table 11-1. p-values for predicted kinetics among treatment conditions compared against the 5 week Normal and 5 week MD animals.** p-values are presented for each cortical area (columns) and for the predicted kinetics of each receptor type(rows). Cortical areas are broken up into comparisons against normal (left) and MD (right). When a curve fit was applied, the equation, degrees of freedom (df), R2 value and exact p-value are listed.
